# Abundance and expression of *hgcAB* genes and mercury availability jointly explain methylmercury formation in stratified brackish waters

**DOI:** 10.1101/2022.02.08.479533

**Authors:** Eric Capo, Caiyan Feng, Andrea G. Bravo, Stefan Bertilsson, Anne L. Soerensen, Jarone Pinhassi, Moritz Buck, Camilla Karlsson, Jeffrey Hawkes, Erik Björn

## Abstract

Neurotoxic methylmercury (MeHg) is formed by microbial methylation of inorganic divalent Hg (Hg^II^) and constitutes severe environmental and human health risks. The methylation is enabled by *hgcA* and *hgcB* genes, but it is not known if the associated molecular-level processes are rate-limiting or enable accurate prediction of MeHg formation in nature. In this study, we investigated the relationships between *hgcA* genes and MeHg across redox stratified water columns in the brackish Baltic Sea. We found that the abundance of *hgcA* genes and transcripts combined with the concentration of dissolved Hg^II^-sulfide species were strong predictors of both Hg^II^ methylation rate and MeHg concentration, implying their roles as principal joint drivers of MeHg formation in these systems. In establishing relationships between *hgcA* genes and MeHg, we advance the fundamental understanding of mechanistic principles governing MeHg formation in nature and enable refined predictions of MeHg levels in coastal seas in response to the accelerating spread of oxygen deficient zones.

## Introduction

The broad dispersal and local accumulation of mercury (Hg) causes large environmental, socio-economic and public health impacts [1]. Future environmental Hg concentrations and human exposures depend on the success of efforts to reduce anthropogenic Hg emissions by implementing The Minamata Convention on Mercury [2] and the impact of ecosystem processes on the reactivity and mobility of legacy Hg [3–5]. The biological formation of methylmercury (MeHg) is a key process as this Hg species is neurotoxic and bioaccumulates in aquatic food webs [6], leading to enhanced exposure of Hg to wildlife and humans [1, 7].

In the environment, MeHg is predominantly formed by methylation of inorganic divalent Hg (Hg^II^) and is mediated by microorganisms carrying *hgc* genes (*hgcA* and *hgcB*) coding for a corrinoid protein and a ferredoxin [8]. Mercury methylating microorganisms thrive in oxygen-deficient environments (e.g., rice paddies, wetlands, sediments, anoxic waters) in which redox conditions play an important regulating role for both the activity of Hg^II^ methylating microorganisms and the availability of Hg^II^ for methylation [9]. Beyond sulfate-reducing bacteria [10], putative or confirmed Hg^II^ methylators have more recently been reported among iron-reducing bacteria, methanogens, fermenters and syntrophs [11–14].

The expression of *hgc* genes is a prerequisite for MeHg formation [8], but no study has demonstrated a quantitative relationship between the abundance of *hgc* genes or transcripts and the amount or formation rate of MeHg. It is thus uncertain if rates of MeHg formation are constrained by the molecular-level methylation processes mediated by the *hgc* genes. Previous studies have shown that the ability of microorganisms to methylate Hg^II^ can be constitutive [15–17] and that Hg exposure not necessarily triggers *hgc* gene expression [15]. Moreover, uncertainties remain about the identity and metabolism of Hg^II^-methylating microorganisms and the specific, and likely variable, environmental drivers that determine their distribution and activity. Accordingly, refined, in-depth sequencing and quantification of *hgc* genes and transcripts as well as other functional genes potentially supporting the Hg^II^ methylation process may provide new information about the distribution and activity of Hg^II^-methylating microorganisms and be important for resolving their roles in Hg^II^ methylation in the environment.

Oxygen deficiency is spreading in coastal seas and the global ocean because of anthropogenic eutrophication and global warming [18, 19]. As one example, the Baltic Sea has seen a dramatic expansion of hypoxic-anoxic water zones over the past 30 years and today holds one of the largest oxygen deficient sediment areas in the world [20, 21]. The expansion of redox-stratified water columns potentially opens new habitats for Hg^II^ methylators and shifts the availability of Hg^II^ to such microorganisms. Higher MeHg concentrations and molar ratios of MeHg to total Hg (HgT) have generally been observed in oxygen depleted waters of redox-stratified coastal seas but contrasting vertical MeHg profiles in anoxic/euxinic zones have been reported. Notably, MeHg concentration maxima have been observed at the hypoxic-anoxic interface [22–24] or in deeper euxinic water layers [23–26]. The reasons for these differences in profiles are not clear and there are still significant uncertainties regarding factors and processes that control MeHg formation in oxygen depleted coastal waters.

Here, we investigate the relationship between the abundance of *hgc* genes and their expression (i.e., *hgc* transcripts), Hg^II^ chemical speciation (controlling Hg^II^ availability), Hg^II^ methylation rate constants and MeHg concentrations measured across vertical profiles of the water column at the Landsort Deep and the Gotland Deep in the central Baltic Sea. Both sites have a redox stratified water column with oxygenated surface water and high dissolved sulfide and MeHg concentrations below the redox transition zone [26]. We test the hypothesis that *hgc* gene abundance or expression controls Hg^II^ methylation rate and accordingly also MeHg concentrations across the redox stratified water column. Our study contributes to a better understanding of MeHg formation in nature and enables refined predictions of expected future changes in MeHg levels in coastal seas following the spread of oxygen deficiency.

## Results and Discussion

### Contrasting redox transition zones at the Landsort Deep and the Gotland Deep

In the central Baltic Sea, pelagic redoxclines are present in several locations, including at the Landsort Deep and Gotland Deep where we collected water samples in August 2019 at stations BY32 and BY15, respectively (Fig 1). Thermoclines were present at 15-40 m water depths for both stations and haloclines at 50-90 m and 60-150 m at BY32 and BY15 stations, respectively (Fig 2A&D, Datasheet 1A). Normoxic water layers (defined as O_2_ concentrations > 2 mL L^-1^) were present from the surface to ~60 and ~70 m depth at the BY32 and BY15 stations, respectively. At BY32, the redox transition zone (here defined as water with O_2_ concentrations between 0.1- and 2-mL L^-1^) was sharp and distinct from ~70 to 80 m. In contrast, the redox transition zone was more extended and distorted reaching from ~80 to 150 m at BY15. The reason for this difference between the two stations is turbulent mixing with oxygenated saline water inputs from the North Sea moving below the halocline to the Gotland basin. These processes substantially impact the redoxcline [26, 27] in the Gotland Deep (BY15 station) but less so in the Landsort Deep (BY32 station; Fig. 1). Correspondingly, euxinic conditions (defined as water with <0.1 mL L^-1^ O_2_ and detectable H_2_S) appeared already at ≤ 80 m at BY32 but at ~150 m at BY15. At greater depths, H_2_S concentrations exceeded 30 μM at both stations and reached a maximum of 100 μM below 200 m at the BY15 station.

**Figure 1:**
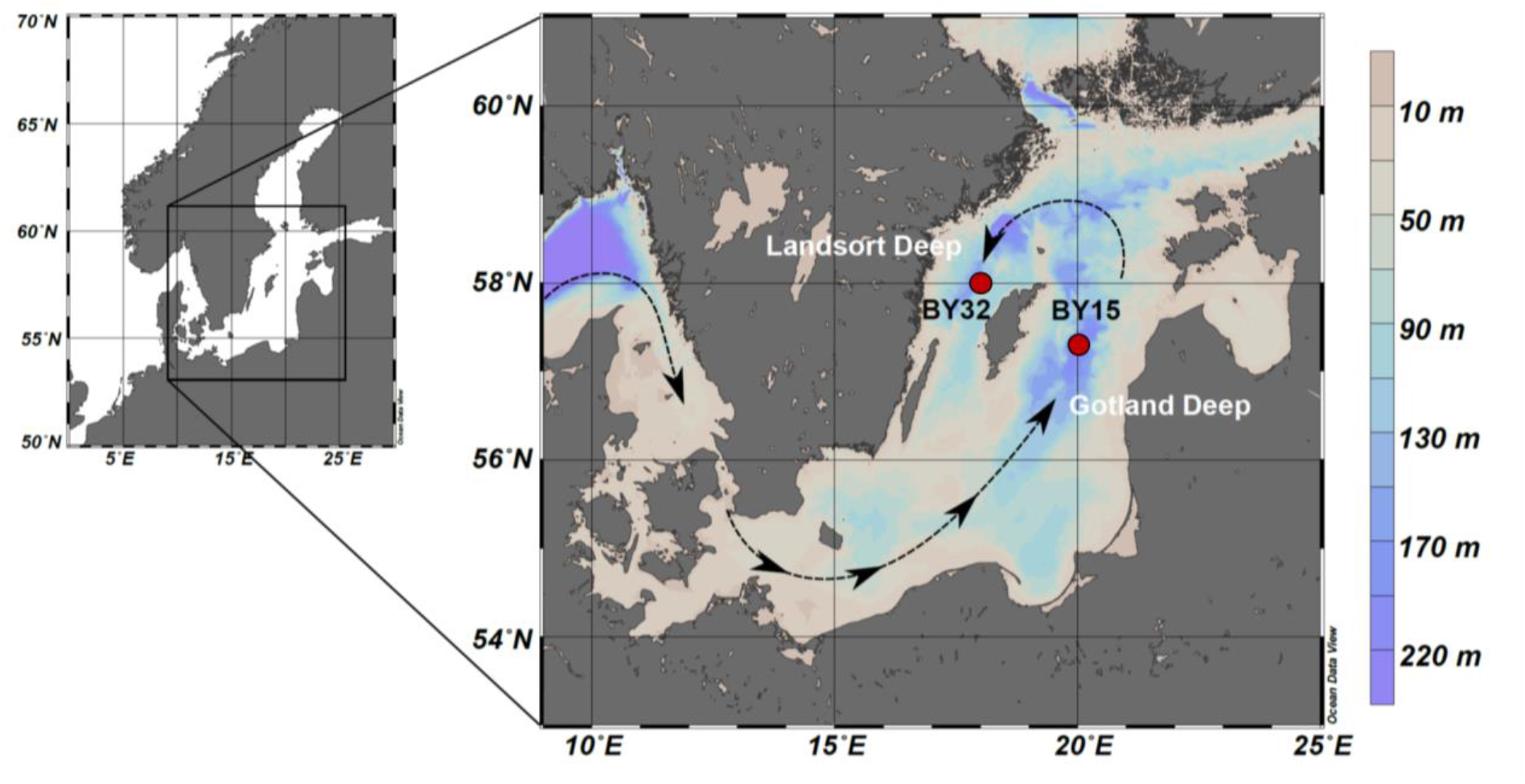
Location of the stations BY32 in the Landsort Deep and BY15 in the Gotland Deep in the central Baltic Sea. Black arrows show the entry point and primary circulation of saline North Sea water below the halocline relevant for this study [28].

**Figure 2.**
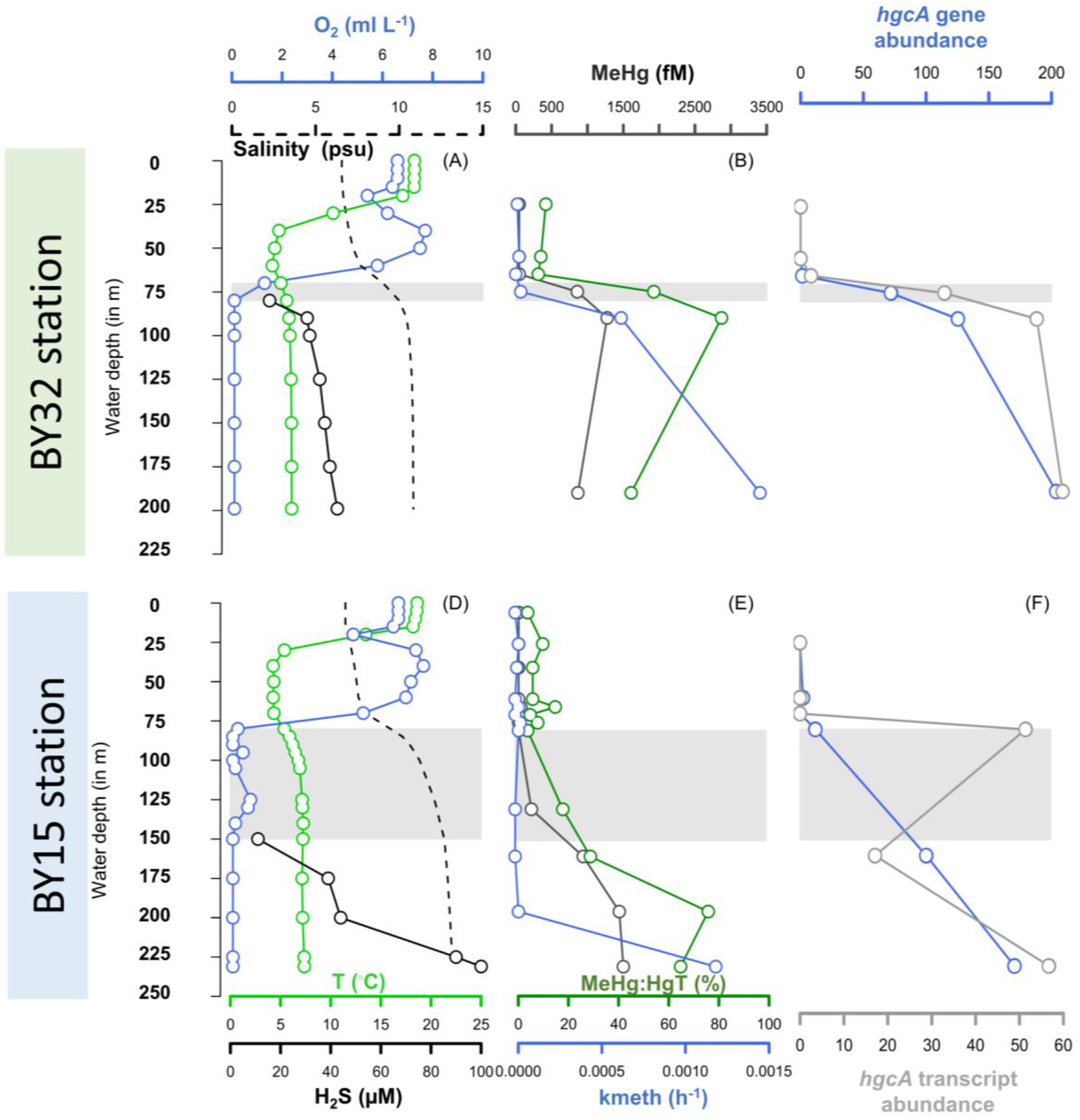
Water depths profiles of ancillary parameters (temperatures, salinity, oxygen, hydrogen sulfide) (A) (D), MeHg concentrations, MeHg/HgT molar ratio (%) and Hg^II^ methylation rate constant (*k*_meth_) in unfiltered water samples (B) (E) and *hgcA* gene and transcript abundance (coverage values in reads/bp normalized with coverage values from the housekeeping gene *gyrB* (D) (F) for stations BY32 (panels A-C) and BY15 (panels D-F).

### Methylmercury concentration and formation rate in relation to redox condition

The vertical distributions of HgT and MeHg concentrations and the molar ratio of MeHg/HgT measured in the water samples increased in the order: normoxic < redox transition < euxinic water zones (Fig. 2B&E, Datasheet 1A). While the HgT concentration increased by a factor of ~4 from normoxic to euxinic waters (from 290-640 to 1800-3000 fM), the increase in MeHg concentration was considerably higher with a factor of ~23 (from <50 to 1000-1300 fM). The MeHg/HgT molar ratio increased accordingly and was <17%, 21-60% and 50-90% in the normoxic, redox transition and euxinic water zones, respectively. The vertical profiles in these MeHg parameters closely followed the respective redox stratification profile at the two stations (described by the O_2_ and H_2_S concentrations) and thus increased sharply over a comparably narrow depth interval of ~20 m at BY32 whereas the increases were more gradual and extended over a ~100 m depth interval at BY15.

The Hg^II^ methylation rate constant (*k*_meth_, determined in incubation assays with an isotopically enriched ^199^Hg^II^ tracer) followed the same overall vertical distribution across the three redox zones as MeHg concentration and MeHg/HgT molar ratio, with highest values in the euxinic zone. The large increase in the euxinic zone, however, occurred at greater depths and higher H_2_S concentration for *k*_meth_ as compared to MeHg concentration and MeHg/HgT ratio at both stations (Fig. 2, Datasheet 1A). The *k*_meth_ value was not detectable (<0.030×10^-3^ h^-1^) in normoxic water samples but increased continuously throughout the euxinic zone and followed the H_2_S concentration reasonably well (Fig 2). At BY32, the *k*_meth_ value was close to detection limit (0.032×10^-3^ h^-1^) in the lower part of the redox transition zone (75 m depth) and then increased substantially with depth and H_2_S concentration in the euxinic zone (0.63×10^-3^ h^-1^ and 1.47×10^-3^ h^-1^ at 90 and 190 m, respectively). At BY15, the *k*_meth_ value was below detection limit throughout the water column until the largest depth in the euxinic zone where a high *k*_meth_ value was observed (1.20×10^-3^ h^-1^ at 230 m). MeHg demethylation rate constants (*k*_demeth_) for samples incubated in the dark were detectable only for the greatest depth at each station (7.6×10^-3^ h^-1^ at BY32, 190 m and 5.1×10^-3^ h^-1^ at BY15, 230 m, Datasheet 1A). Combined, these results suggest that MeHg concentration and MeHg/HgT ratio in the oxygen deficient water zones are mainly controlled by *in situ* Hg^II^ methylation except at the deepest euxinic water where also MeHg demethylation was high.

Overall, the HgT and MeHg concentrations and MeHg/HgT molar ratio observed in our study were in the same ranges as previous observations for redox-stratified waters in the Baltic Sea [23, 26], although Soerensen et al. [26] observed maximum HgT concentrations that exceeded the levels measured in our study. Differences in depth of maximum MeHg concentration and/or MeHg/HgT ratio have been reported among or within studies on redox stratified coastal systems, with maxima located either at the interface between the redox transition (or hypoxic) zone and the euxinic (or anoxic) zone [22–24, 29] or in deeper euxinic water layers with H_2_S concentrations in the μM range [23–26]. Notably, the MeHg concentration and MeHg/HgT ratio, which are the parameters normally determined in studies on MeHg in coastal seas, are proxies for net formation of MeHg [30]. These parameters thus represent the net result of Hg^II^ methylation and MeHg demethylation (and are in some cases also influenced by input and output of Hg^II^ and MeHg in the studied system) and do not enable to study the two processes separately. There are very few studies determining the individual rate constants *k*_meth_ and *k*_demeth_ in redox stratified coastal seas [26]. Our study demonstrates that MeHg gross formation (i.e., methylation rate of Hg^II^) is largest in euxinic waters with high H_2_S concentration, rather than at the interface between the redox transition and euxinic zones. These results highlight the value of determining rate constants for Hg^II^ methylation and MeHg demethylation along with MeHg concentration and MeHg/HgT ratio to fully characterize MeHg cycling in coastal seas.

### Chemical speciation modelling revealed the highest availability of Hg^II^ for methylation in the euxinic zone

The chemical speciation of Hg^II^ in the different pelagic redox zones in the Central Baltic Sea have previously been described conceptually and by thermodynamic modelling [26]. We used the same thermodynamic model to verify the Hg^II^ speciation for the conditions at the time of our sampling cruises (Supplementary Information, Table S1) but improved the accuracy by including DOC observations from the cruise rather than the generic values used in Soerensen et al. [26]. The results of the model suggest that in normoxic waters, Hg^II^ is present as complexes with thiol groups associated with dissolved or particulate OM. In the redox transition zone, the model predicts that Hg^II^ complexes with thiols dominate at H_2_S concentrations lower than ~0.003 μM. These Hg^II^-thiol complexes are expected to predominantly be associated with particulate aggregates consisting of Fe^III^-oxyhydroxides and OM in the transition zone [22, 26, 31]. The solid-phase metacinnabar, HgS(s), dominates the Hg^II^ speciation at H_2_S concentrations in the fairly narrow range (in absolute terms) of 0.003 to 0.25 μM, whereas dissolved Hg^II^-sulfide species outcompete HgS(s) at H_2_S concentrations exceeding 0.25 μM. The distribution among dissolved Hg^II^-sulfide species are 86.5% HgS2H^-^, 10.1% Hg(SH)2 and 3.4% HgS2^2-^ at pH 7.3, as used in our model, irrespective of H_2_S concentration above 0.25 μM.

It is important to note that the particulate Hg^II^ phases present in the redox transition zone have low availability for microbial Hg^II^ methylation compared to the dissolved Hg^II^-sulfide species dominating in the euxinic water zone [32, 33]. Previous studies proposing maximum MeHg formation at the interface of redox zones based on MeHg concentration or MeHg/HgT ratio (i.e., proxies for MeHg net formation) have explained this observation with a decreased Hg^II^ availability for methylation at high μM concentrations of H_2_S. These conclusions, however, were partly based on inaccurate speciation modeling of Hg^II^ with models largely influenced by the species HgS^0^(aq) [34], which has been refuted in later studies [35, 36], and/or based on speciation modelling of soils and sediment with orders of magnitude higher Hg^II^ concentrations. In contrast, our speciation modeling predicted highest availability for methylation of Hg^II^ in the euxinic zone, at H_2_S concentrations exceeding 0.25 μM, through the formation of dissolved Hg-sulfide species. This result is in line with highest observed *k*_meth_, MeHg concentration and MeHg/HgT ratio in the euxinic zone in our work.

### Distribution and activity of Hg^II^-methylating taxa is tightly linked to redox conditions

Previous metagenomic studies have demonstrated the presence of *hgc* genes in oxygenated marine pelagic systems, suggesting a potentially important role of certain aerobic lineages, e.g., *Nitrospina*, in marine Hg^II^ methylation [37–40]. So far, less attention has been given to oxygen-deficient marine environments [40–42] where other Hg^II^ methylating lineages are expected to thrive. Consistent with other recent findings [40], our results based on the sequencing of metagenomes and metatranscriptomes (Datasheet 1B) from water samples demonstrated that redox conditions is a major factor controlling the presence and abundance of potential Hg^II^ methylators in the central Baltic Sea water column (Fig. 2&3). An overview of the link between redox conditions, and the composition and metabolism of the overall microbial community is described in Supplementary Information (including Fig. S1).

**Figure 3:**
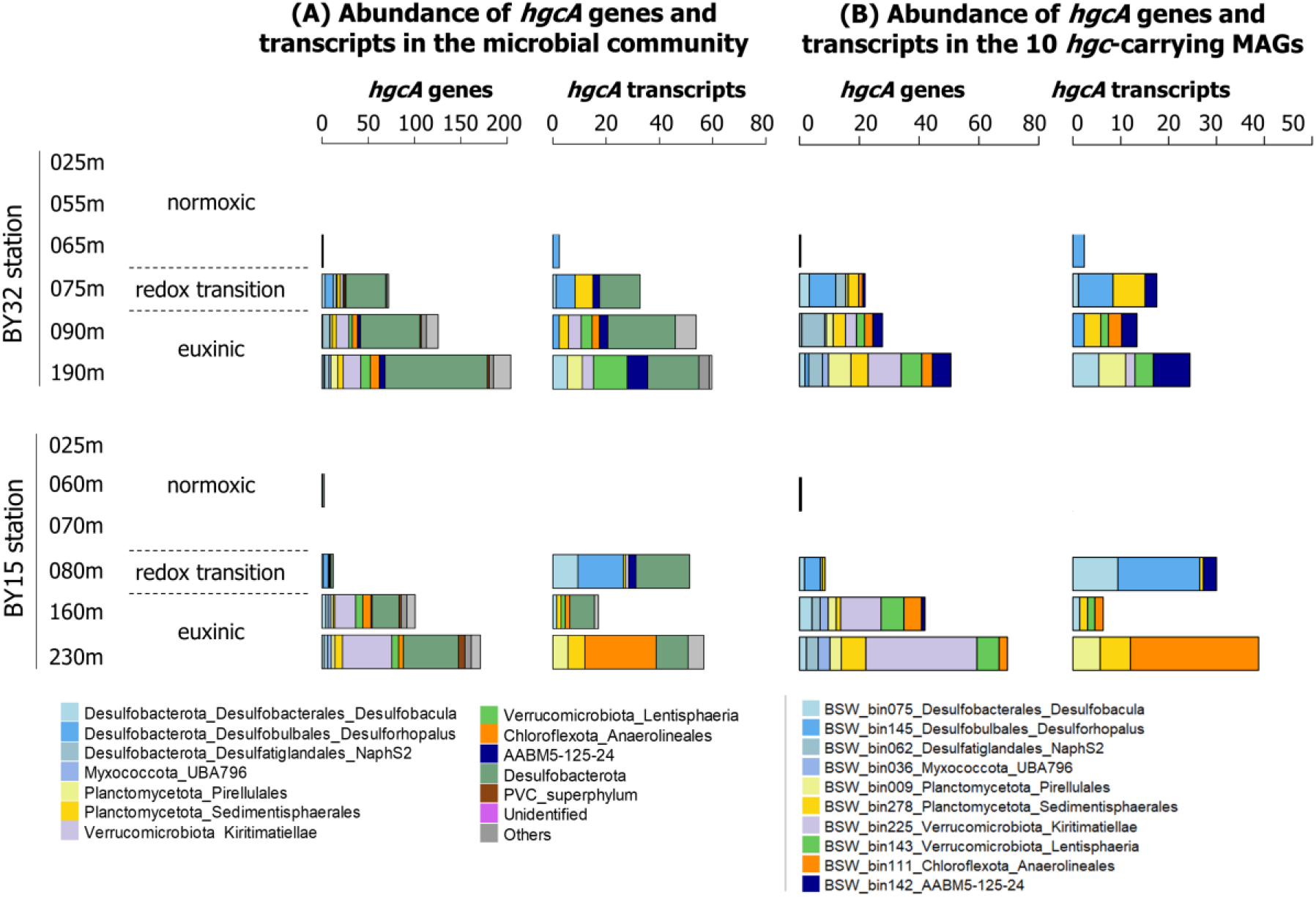
Distribution of *hgcA* genes in terms of normalized abundance (counts) of both total genes and transcripts for all *hgcA* genes detected in the water metagenomes (A) and the 10 *hgcA*-carrying MAGs (B). Color codes correspond to the taxonomic identification of each microbial group, see bottom panels for correspondence.

We focused here on the distribution and activity of microorganisms carrying and expressing *hgc* genes, and thus, potentially involved in Hg^II^ methylation. In terms of abundance (i.e., normalized coverage values as described in Material and Methods section), the overall pattern was the same at both stations. The *hgcA* (Fig. 3A, Datasheet 1C) and *hgcB* (Fig. S2, Datasheet 1C) genes and their transcripts were either absent or at very low levels in the normoxic zone, increased in the redox transition zone and were present in highest abundances in the euxinic zone. One exception to this general pattern was the lower abundance of *hgc* transcripts at BY15 160 m as compared to other samples from the euxinic zone (Fig. 3A, Fig. S2). Analogously to the redox and Hg parameters, the increase in *hgc* gene and transcript abundances across the redox zones were sharper at the BY32 compared to BY15 station. At both stations, *hgc* transcripts from members of Desulfobacterota and PVC superphylum (i.e., Planctomycetota, Verrucomicrobiota) dominated in both the redox transition and euxinic zones, while *hgc* transcripts from Chloroflexota and AABM5-125-24 were only abundant in euxinic waters samples (Fig. 3A, Fig S2).

The reconstruction of 127 metagenome-assembled-genomes (MAGs; Datasheet S1E) revealed that 10 of them carried *hgc* genes that contributed to, respectively, 44 and 59 % of the *hgcA and hgcB* transcripts found in the euxinic zone from both stations (Datasheet 1C). A detailed characterization of these 10 putative Hg^II^ methylators and their metabolic traits can inform about the metabolic processes that drive Hg^II^ methylation and about environmental factors that control their distribution and activity [9]. To investigate the expression levels of *hgc* genes found in MAGs, we compared the expression level of *hgc* and metabolic capacity genes to expression levels of the housekeeping gene *gyrB* detected in the 10 *hgc*-carrying MAGs (Fig. 4). For each MAG, expressions levels were considered as high if they surpass *gyrB* expression level by 0.005 (in coverage values of transcripts). We found that *hgc* genes were expressed in the redox transition and euxinic zones by MAGs featuring in total five different metabolic capacities. Sulfate reduction, fermentation and hydrogen oxidation were predominant while the contributions from aerobic and micro-aerophilic respiration processes were smaller (Fig. 4, Datasheet 1F).

**Figure 4.**
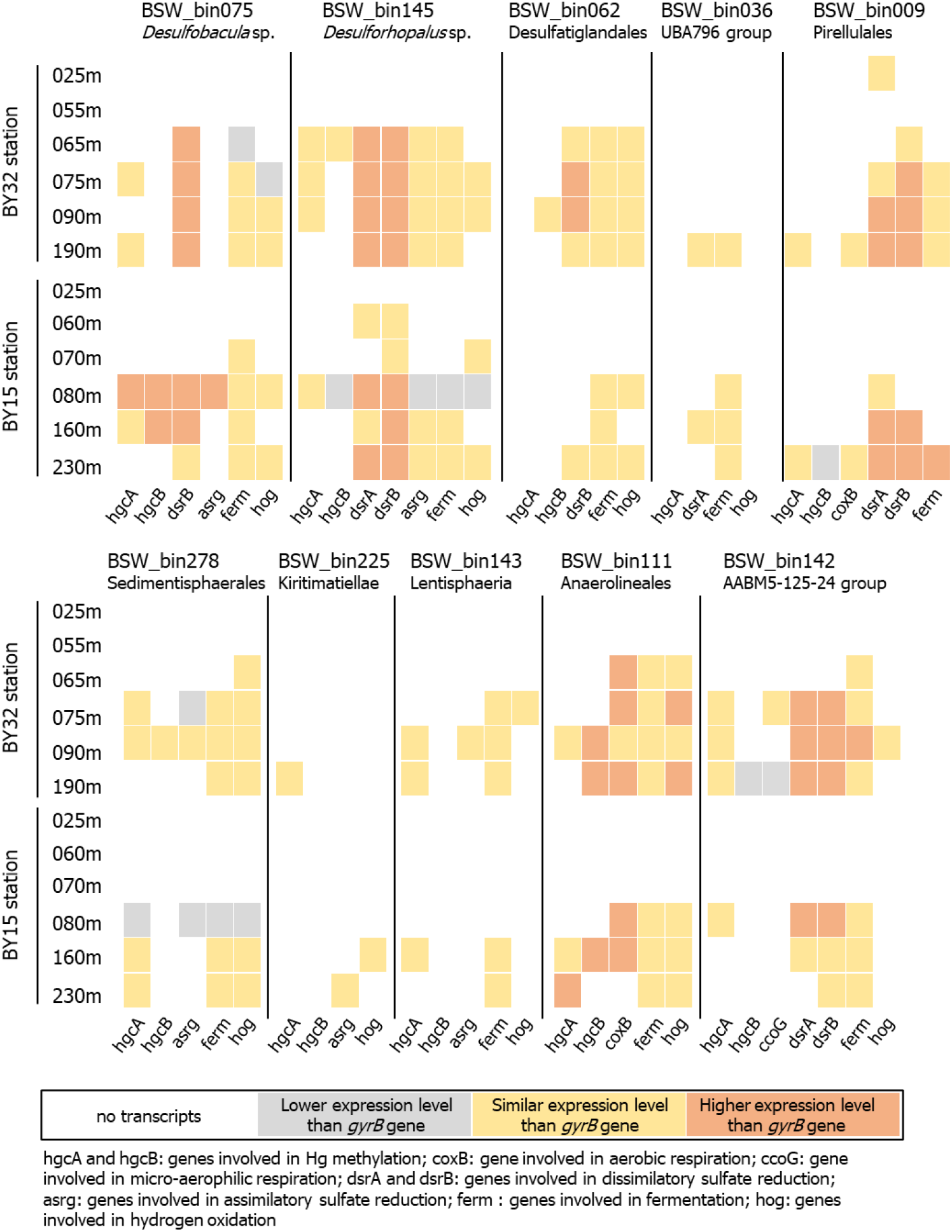
Expression profiles of functional genes by *hgc*-carrying MAGs found in BY32 and BY15 stations. For each MAG, expression levels were calculated comparing the coverage values of transcripts from functional genes to the coverage values of the transcripts from the housekeeping gene *gyrB*. Colors denote expression levels of functional genes lower (gray), similar (yellow) and higher (orange) than the expression level of *gyrB* genes. Expression levels were considered at similar if the differences between coverage values were less than 0.005 (see Datasheet 1G for exact values).

Two Desulfobacterota MAGs - *bin145* (*Desulforhopalus* sp.) and *bin075 (Desulfobacula* sp.) - expressed *hgc* genes in the redox transition and euxinic zones (BY32 75-90 m, BY15 80-160m m water depth) while *bin062* (Desulfatiglandales NaphS2 group) expressed *hgcB* transcripts (but not *hgcA*) only at BY32 90 m (Fig. 3B, Fig. 4). Notably, the *hgc* expression level from *bin075* was high in the redox transition zone of BY15 station (Fig. 4). These three MAGs also expressed genes involved in sulfate reduction, fermentation and hydrogen oxidation, which are known metabolic capacities for Desulfobacterota members [43–45]. For genes involved in sulfate reduction, including *dsrAB* genes, expression levels of these three MAGs were found at similar or higher expressions levels than *gyrB;* see for instance the strong expression of *bin0145* in the redox transition and euxinic zones at BY32 station (Fig. 4). The fourth Desulfobacterota MAG, *bin036* (Myxococcota UBA796 group) did not express any *hgc* genes but expressed sulfate reduction and fermentation genes at similar level as the expression of the *gyrB* gene. Two MAGs from the PVC superphylum expressed *hgc* genes in both the redox transition zone and euxinic zone, with a dominance of the Planctomycetes *bin278* (Sedimentisphaerales) in BY32 75-90 m and BY15 160-230 m and a dominance of the *bin009* (Pirellulales) in the deepest water samples at both stations (BY32 190 m, BY15 230 m) (Fig. 3B). In accordance with recent findings [46], *bin009* expressed high levels of sulfate reduction genes in euxinic samples (Fig. 4). The two *hgc*-carrying Verrucomicrobia MAGs, that are *bin225* (Kiritimatiellae) and *bin143* (Lentisphaeria) expressed high levels *hgcA* genes in euxinic samples (Fig. 4) with the highest levels at BY32 190 m for *bin143* (Fig. 3B). The single Chloroflexota representative (*bin111*, Anaerolineales) expressed *hgc* genes only in the euxinic zone (Fig. 3B) in which it also produced high expression levels of transcripts involved in hydrogen oxidation and, surprisingly, transcripts involved in aerobic respiration (Fig. 4). Although *bin142* (AABM45-125-24 phylum) expressed *hgc* transcripts in the redox transition zone at both stations, they were expressed in the euxinic zone only at BY32. This MAG also expressed high levels of genes involved in sulfate reduction and fermentation at the same locations, which is in accordance with the limited knowledge so far about the ecology of members affiliated with this phylum [47]. Overall, taxonomical differences were found in the *hgc*-carrying MAGs populating the redox transitions and euxinic zones but no clear differences were found in their metabolic capacities. These observations support the hypothesis that Hg methylators exhibit versatile metabolic capacities [42, 48, 49] and that microbial Hg methylation cannot be associate to a single metabolism.

### Relationships between *hgc* abundances, Hg^II^ speciation and MeHg formation

We investigated potential relationships between gene and transcript abundances of *hgc* and the three Hg parameters *k*_meth_, MeHg concentration and MeHg/HgT molar ratio. There was a correlation between *hgcA* and *hgcB* gene abundances (r^2^=0.95, p < 0.001) and transcripts abundances (r^2^=0.95, p < 0.001) and at the gene level the correlations with Hg parameters were significant for both *hgcA* or *hgcB* (Table S3). At the transcript level, however, only *hgcA* consistently showed significant correlations with all Hg parameters and we therefore focus on *hgcA* in the further discussion Significant positive Spearman rank correlations were found between *k*_meth_ values and *hgcA* gene (r^2^=0.58, p = 0.048) and transcript abundance (r^2^=0.65, p = 0.023) detected in the water layers from both stations (Fig. 5, Table S2). Low *k*_meth_ (<0.04×10^-3^ h^-1^) was found in the redox transition samples and in one euxinic sample (n=3) while high *k*_meth_ (>0.6×10^-3^ h^-1^) was found in the other euxinic samples (n=3) (Fig. 5). These two groups (n=3 each) contained samples from both stations showing that the grouping in samples with low and high *k*_meth_ was not site specific. The highest *k*_meth_ was observed for the euxinic samples with highest *hgcA* gene abundance and expression (Fig. 5). We propose that this apparent “threshold” in the relationship between *k*_meth_ and *hgcA* gene or transcript abundances was primarily explained by the chemical speciation of Hg^II^. As discussed above, dissolved Hg^II^-sulfide complexes, which have a comparably high availability for methylation, dominated the Hg^II^ speciation in euxinic samples (black colored data points in Fig. 5). In contrast, adsorbed phases of Hg^II^ with Fe^III^-OM aggregates [22, 31] and the solid-phase HgS(s), with much lower availability for methylation [32, 33], are expected to dominate the speciation in the redox transition zone (yellow colored data points in Fig. 5).

**Figure 5.**
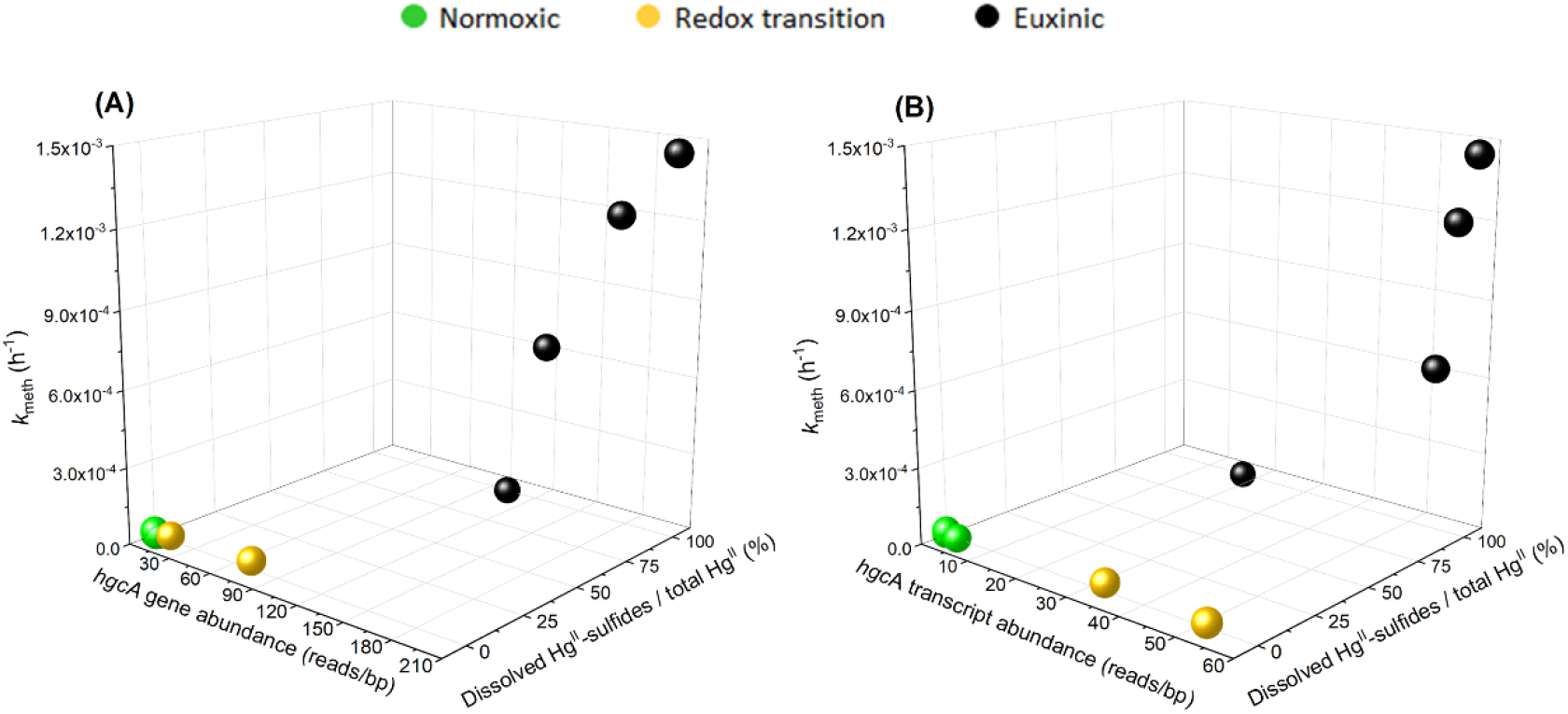
Joint relationship between *hgcA* gene (A) or transcript (B) abundances, fraction (% of total Hg^II^) of dissolved Hg^II^-sulfide species and *k*_meth_. Black colored data points correspond to samples from the euxinic zone, with a Hg^II^ speciation dominated by dissolved Hg^II^-sulfide complexes, and yellow and green data points represent samples from the redox transition and normoxic zones, respectively, both void of such complexes.

We took into account both *hgcA* abundances and Hg^II^ availability for MeHg formation (Fig. 5) by partial least squares projection (PLS) modeling. Two separate PLS models with the concentration of dissolved Hg^II^-sulfide species and either the *hgcA* gene (r^2^=0.83, q^2^=0.62) or transcript (r^2^=0.69, q^2^=0.59) abundance as X-variables were generated (Supplementary Information, Table S3). Compared to the models presented above, with *hgcA* abundances only, the fit was improved when taking into account also Hg^II^-sulfide species in the models. The results support the hypothesis that Hg^II^ methylation across the three pelagic redox zones is controlled jointly by the abundance of *hgcA* and the concentration of dissolved Hg^II^-sulfide complexes (Fig. 5). These two drivers of MeHg formation followed partly different vertical profiles across the redox zones. The *hgcA* genes and transcripts were consistently present and expressed both in the redox transition and euxinic zones, showing that there is a high microbial potential for Hg^II^ methylation in both zones. In the transition zone, however, the rate of Hg^II^ methylation was below or close to detection limit due to the low availability for methylation of Hg^II^, which most likely was rate limiting for MeHg formation in this zone. In euxinic waters, the availability of Hg^II^ was high throughout the entire zone due to formation of dissolved Hg^II^-sulfide species and the Hg^II^ methylation rate was limited by the abundance of *hgcA* genes/transcripts.

One euxinic sample (BY15 160m) exhibited non-detectable *k*_meth_ (Fig. 5). The composition and metabolism of the overall prokaryotic community (Fig. S1) in this sample, as well as the composition of Hg^II^-methylating groups (Fig. 3A), were consistent with other samples from the euxinic waters. However, compared to the samples with high *k*_meth_, the total abundance of *hgcA* transcripts were lower in this sample (Fig. 3A, Fig S2) and we suggest this might explain the low *k*_meth_ values measured for this sample. The few detected *hgcA* transcripts were associated with four MAGs, including, a *Desulfobacula* MAG (*bin075*) and a Anaerolineales MAG (*bin111*). Interestingly, these *hgc*-expressing MAGs contributed less, in proportion, to the expression of genes involved in the metabolic capacities of the overall community (i.e., transcript expression levels for sulfate reduction and fermentation respectively) in this sample (0.29 and 0.01 %) compared to other euxinic samples (0.78-4.01, 0.03-0.04 %) (Datasheet 1D, 1F). This implies that the *hgc*-carrying MAGs could be outcompeted by non-*hgc*-carrying MAGs in this layer leading to reduced capabilities of the resident community to methylate Hg^II^.

We evaluated if other processes could explain the apparent threshold in the relationship between *k*_meth_ and *hgcA* genes and transcripts. We considered the central metabolic capacities of microorganisms expressing *hgc* genes and the composition of organic matter (OM) in the water columns. Laboratory experiments with isolated microorganisms have demonstrated large differences in Hg methylation capacity within and between microbial phyla [50], but it is still not clear if this process is linked to their essential metabolic capacities. Our results showed that there was no systematic difference in metabolic capacities of *hgcA* expressing MAGs in the redox transition and euxinic zones (Fig. 4) that could explain the large differences in observed *k*_meth_. Previous studies have demonstrated that the origin and degradation status of OM can control Hg^II^ methylation in contrasting lakes [51] and boreal ponds [52]. These previous studies used GC-MS and fluorescence spectroscopy (respectively). In our study, we characterized the molecular composition of the dissolved OM (DOM) in all samples using liquid chromatography – electrospray ionization mass spectrometry [53]. The data obtained show that the DOM composition did not vary within the euxinic zone and differed only marginally between the redox transition and euxinic zones (Supplementary information). Hence, for these water columns it is unlikely that the composition of DOM was an important contributing factor to the observed differences in *k*_meth_.

The *hgcA* gene and transcript abundances were significantly correlated also with the ambient MeHg concentration (r^2^=0.87, p<0.001 and r^2^=0.78, p<0.001, respectively) and the MeHg/HgT molar ratio (r^2^=0.63, p=0.027 and r^2^=0.58, p=0.048, respectively) (Table S2). Further, like the results for *k*_meth_, PLS models considering both *hgcA* gene or transcript abundance and concentration of dissolved Hg^II^-sulfide complexes gave a good fit for both MeHg concentration and the MeHg/HgT molar ratio (r^2^=0.82, q^2^=0.70 and r^2^=0.67, q^2^=0.40, respectively for PLS models with *hgcA* gene abundance and Hg^II^-sulfide complex concentration). In principle, MeHg concentration and MeHg/HgT ratio are not only determined by *hgcA*-driven Hg^II^ methylation but also by the MeHg demethylation rate and MeHg and Hg^II^ fluxes. It is thus noteworthy that the *hgcA* gene and transcript abundances, together with the concentration of dissolved Hg^II^-sulfide species, could explain most of the variance both in *k*_meth_ determined from an added Hg^II^ isotope tracer in incubation experiments and in ambient MeHg concentration and MeHg/HgT in the water samples. The tight link between all the parameters related to MeHg formation shows the importance of methylation of Hg^II^, driven by *hgcA* gene and transcript abundances and concentration of dissolved Hg^II^-sulfide species, for build-up of the ambient MeHg pool in these redox stratified waters.

### Environmental implications

The presence of *hgc* genes has proven to be a reliable predictor of Hg^II^ methylation capability in microorganisms [50, 54]. Indeed, deletion of the *hgc* genes impairs Hg^II^ methylation capacity [8]. Further, Qian et al. [55] observed a relationship between HgcA protein concentration and amount of formed MeHg in laboratory assays with a sulfate-reducing bacterium. However, no study has demonstrated a quantitative relationship between the abundance of *hgc* genes or transcripts and the amount or formation rate of MeHg, neither in environmental samples nor in laboratory culture experiments [15–17, 56]. It has thus remained uncertain if rates of MeHg formation are constrained by the molecular-level methylation processes mediated by the *hgc* genes. In general, a positive relationship between the abundance of gene or transcripts and corresponding process rates is often assumed but rarely demonstrated in natural systems, mainly because of a lack of fundamental understanding on the factors that control the activity of the microorganisms and thus the gene transcription [57].

In this study, we show significant relationships between *hgcA* gene abundance as well as *hgcA* gene transcripts and all three quantified MeHg parameters *k*_meth_, MeHg/HgT molar ratio and MeHg concentration. Our results further show that models with both *hgcA* gene or transcript abundances and Hg^II^ chemical speciation improved the prediction of MeHg formation in redox stratified pelagic waters, suggesting that *hgcA* and Hg^II^ availability jointly control MeHg formation in such environments. These key findings are conceptually illustrated in Fig. 6. The observed presence of *hgcA* genes and transcripts in the redox transition zone indicates that there is microbial potential for MeHg formation under such conditions. The chemical speciation and availability for methylation of Hg^II^ is however expected to fluctuate substantially even with moderate variations in H_2_S concentration in this zone. As a consequence, large spatial and temporal variability in MeHg formation, driven by H_2_S concentration, may be expected. In contrast, Hg^II^ availability for methylation is high throughout the euxinic water zone and the MeHg formation is instead controlled by the abundance and expression of *hgcA* genes.

**Figure 6.**
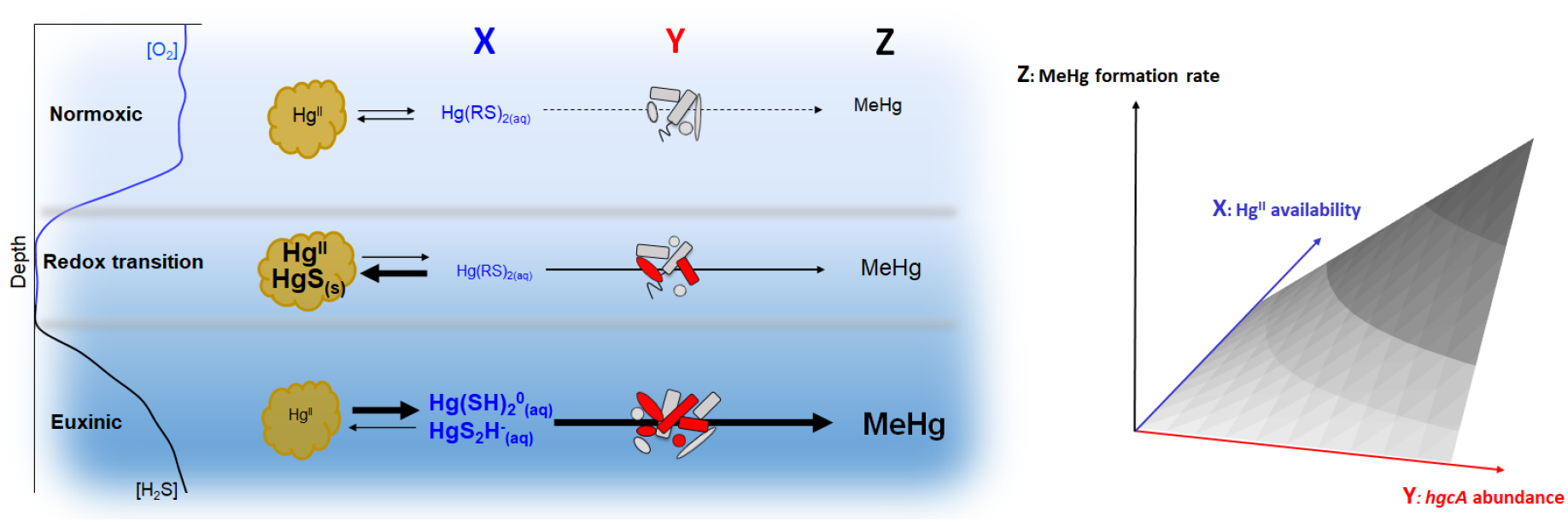
Conceptual illustration of *hgcA* genes and Hg^II^ availability jointly controlling MeHg formation in redox stratified brackish waters. The left panel illustrates vertical profiles of O_2_ and H_2_S across the three pelagic redox zones. The center panel illustrates how Hg^II^ availability (X) and *hgcA* gene/transcript abundance (Y) jointly control MeHg formation (Z) across the three redox zones. The Hg^II^ availability is controlled by partitioning between particulate Hg^II^ species (black font), not available for MeHg formation, and dissolved Hg^II^ species (blue font) available for MeHg formation. In euxinic waters, the partitioning is largely shifted to dissolved Hg^II^-sulfide species. Microorganisms not carrying or expressing *hgcA* (grey color) are present throughout the water column while microorganisms expressing *hgcA* (red color) appears in the redox transition zone and increase further in abundance in the euxinic zone. The shift in community composition across the redox zones is illustrated by different shapes of the microorganism cartoons. The conceptual contour plot in the right panel further illustrates how MeHg formation is jointly driven by Hg^II^ availability and *hgcA* gene/transcript abundance.

The native function of the *hgc* genes is still not known [8, 58] but they do not seem to be part of a Hg detoxification system since the gene expression is not induced by Hg exposure and does not confer Hg resistance [15, 57]. Our results suggest that Hg^II^ methylation is, to some extent, associated with the overall microbial activity of certain microorganisms as well as their specific metabolic capacities such as sulfate reduction, fermentation and hydrogen oxidation (Fig. 4A, Figs S2&3). Overall, our results point to these three principal metabolisms among putative Hg^II^ methylators in the redox stratified water column of the central Baltic Sea. We hypothesize that, without having an evident causality link between Hg^II^ methylation and other metabolic processes, the characterization of the biogeochemistry and associated metabolism of microbial communities inhabiting aquatic systems is a novel and useful approach to better predict the presence of Hg^II^ methylation in aquatic systems.

Based on high observed MeHg concentrations in oxygen deficient environments and the fact that the vast majority of known microorganisms capable of Hg^II^ methylation are anaerobes [9], it is expected that current and future spread of oxygen deficiency in coastal seas and the global ocean will increase MeHg formation. However, quantitative predictions have been hampered by a lack of understanding of principle drivers controlling MeHg formation under such conditions. The results of our study benchmark the use of *hgcA* genes and transcripts and their abundances as strong predictors, together with dissolved Hg^II^ sulfide species, for MeHg formation in redox stratified waters. The unraveling of these mechanistic principles governing MeHg formation in oxygen deficient coastal seas is a critical step to refine predictive frameworks for MeHg exposure to marine food webs and humans from current and future spread of oxygen deficiency in coastal waters and the global ocean, as well as for MeHg formation in the environment in general.

## Materials and Methods

### Study sites

Water samples were collected in late August 2019 in the Baltic Sea at station BY32 (57°19′N 20°03′E) in the Landsort Deep and station BY15 (58°01′N 17°58′E) in the Gotland Deep. Temperature, salinity, oxygen, and sulfide were measured on board as a part of the Swedish National Monitoring Program. Samples were analyzed according to HELCOM guidelines (HELCOM Combine, 2014, www.helcom.fi) and the quality-controlled data is available in Datasheet 1A and from the website of Swedish Meteorological and Hydrological Institute (SMHI, 2019, https://sharkweb.smhi.se/hamta-data/). In cases where the determined ancillary parameters did not match our sampling depth, the numerical value of the parameter was determined by linear interpolation from the two nearest depths. Water-sediment interfaces were reached at 204 and 240 m for BY32 and BY15 stations, respectively,

### Water sampling and sample handling

Water samples were collected at six or twelve sampling depths across the redoxcline using 2-L Niskin bottles (attached to a rosette) at the BY32 and BY15 station, respectively. Water from the sampling rosette was collected in 2 L Fluorinated HDPE transfer bottles, and then split into several bottles for different parameters analysis purposes. All the sampling vessels were pre-cleaned according to trace metal protocols [60]. Oxygen deficient water samples from the redox transition and euxinic zones were pushed out of the Niskin bottles with an N2 flow and collected in the transfer bottles inside an N2 filled glove bag to avoid exposure to ambient air. HgT samples were collected in 125 ml amber glass bottles (I-CHEM certified™ 300 series), MeHg and *k*_meth/demeth_ samples were collected in 250 ml Fluorinated HDPE bottles. Samples for HgT and MeHg concentration determination were preserved by addition of 0.1M HCl and stored in the fridge until analysis.

Incubation experiments to determine the rate constants of Hg^II^ methylation (*k*_meth_) and MeHg demethylation (*k*_demeth_) were conducted onboard. Samples were spiked with ^199^Hg^II^ (to 260 pM) and Me^201^Hg (to 2 pM) enriched isotope tracers and incubated in dark at room temperature in an N2-filled glove box and terminated by acidification to 0.1M HCl at time points of 0, 4, 8 (or 12) and 24h.

For DNA and RNA analysis, three 1 L water samples were collected at totally six depths from the normoxic (1 sample), redox transition (2 samples) and euxinic water layers (3 samples) at the two stations. The sampling depths corresponded to that for chemical parameters. All samples were filtered through 0.22 μm sterivex polyethersulfone enclosed filters using a peristaltic pump. After filtration, RNA later^®^ solution was added to each filter and the filters were stored at −20 °C prior DNA and RNA extractions.

### Determination of HgT and MeHg concentrations and *K*_meth_ and *k*_demeth_

The HgT water samples were treated with BrCl and Hg^II^ subsequently reduced to Hg^0^ by SnCl_2_ and stripped off by purging with N2 followed by dual gold amalgamation and Cold Vapour Atomic Fluorescence Spectroscopy (CVAFS) detection (US-EPA Method 1631E). Lab blanks consisting of MQ water (N=6) were lower than LOQ (0.25 pM). Field blanks (N=6) were prepared from MQ water, brought onboard the cruise and treated as the samples and the determined concentration was 0.36±0.07 pM. We subtracted the difference of the field and lab blanks (0.11 pM) from all HgT samples for blank correction. The RSD of triplicate field samples collected from the same water depths was on average of 12%.

MeHg concentrations and *k*_meth/demeth_ rate constant samples were spiked with a Me^200^Hg isotope enriched internal standard in the lab, derivatized with sodium tetraethylborate, purged and trapped on Tenax^®^ adsorbent [61, 62], and subsequently determined by isotope dilution analysis using thermal desorption gas chromatography inductively coupled plasma mass spectrometry (TDGC-ICPMS) measurements. The MeHg sample concentrations were blank corrected by subtracting half the limit of detection (LOD) value, which was 49 fM based on 3 times the standard deviation of low MeHg concentration samples (n = 11). The RSD of triplicate field samples collected from the same water depths was on average of 4%. The *k*_meth_ and *k*_demeth_ rate constants were determined as the slope of linear fits of Me^199^Hg concentration or ln(Me^201^Hg/Me^201^Hgt=0) concentration ratio, respectively, versus time. Rate constants were calculated only for samples with a statistically significant linear regression (p<0.05, Fig. S3) and were considered non-detectable for other samples.

### Hg speciation model

The chemical speciation of Hg^II^ and MeHg across the different water zones was determined by thermodynamic modeling using the WinSGW software from Majo, Umeå, Sweden [63]. We adapted a model developed by Liem-Nguyen et al. [64], which comprised 7 components and 24 species. A full description of the model used in our study is given in Tables S1&S3. As input to the model was used the determined concentrations of Hg^II^ (calculated as HgT-MeHg), MeHg, H_2_S and fixed Cl^-^ (90 mM) and pH (7.3) values, as well as the estimated concentration of thiol functional groups associated with dissolved organic matter (DOM), as 0.15% of the DOC concentration.

### DOM composition

Dissolved organic matter was extracted from water samples according to the method by Dittmar et al. [65]. Sample volumes varied from 350-1000 ml, and DOM extracts were prepared by solid phase extraction so that approximately the same equivalent volume of seawater was analyzed in each case. The methanol extracts were dried down, re-dissolved in 5% methanol in water, and analyzed by LCMS using a Polar C18 column (Phenomenex) with detection by DAD (light absorption), charged aerosol detector (material abundance) and negative mode electrospray ionization mass spectrometry (Orbitrap [66]). To evaluate the DOM composition from the water samples, the intensities throughout the chromatographic separation were summed to produce one peak list for each sample (Fig. S4), to which formulas were assigned (Fig. S5). Additionally, Suwannee River Fulvic Acid (International Humic Substances Society batch 2S101F) was analyzed for quality control before and after the samples. In total, 1840 chemical formulas were found in at least one sample in the dataset. The data were normalized in each sample to sum 1 × 10^6^ and samples were compared pairwise using the Bray-Curtis dissimilarity metric. The dissimilarity matrix was used to perform principal coordinate analysis (Fig. S6).

### Molecular analysis and bioinformatics

DNA and RNA were extracted from one and two filters, respectively, with the FastDNA^®^ SPIN Kit for soil filter and two with the QIAGEN RNeasy mini kit. The TURBO DNA-free kit (Invitrogen) was used to DNase treat extracted RNA and remove DNA contamination and afterwards controlled for residual DNA by a 30-cycle PCR with 16S rDNA primers (27F and 1492R) including milli-Q as negative and DNA from *E. coli* as positive controls. This was followed by bacterial ribosomal RNA depletion using the RiboMinus Transcriptome Isolation Kit (ThermoFisher Scientific). The final quantity and quality of extracted nucleic acids were measured on a NanoDrop One spectrophotometer (ThermoFisher Scientific) and a qubit. Library preparation of DNA and RNA for sequencing was prepared with the ThruPLEX DNA-seq (Rubicon Genomics) and TruSeq RNA Library Prep v2 (Illumina) kits, respectively. The DNA and RNA samples were each sequenced with a paired-end 2 × 150 bp setup at the Science for Life Laboratory (Stockholm) at the SNP&SEQ Technology Platform over an entire Illumina NovaSeq6000 SP4 flow cell.

The bioinformatic procedure is described in detail in the Supporting information. As a brief description of the workflow: (i) the 12 metagenomes were assembled into a co-assembly (ii) reads from both metagenomes and metatranscriptomes were then mapped to the co-assembly (iii) the co-assembly was used to reconstruct metagenome-assembled genomes (MAGs) (iv) protein-coding genes were predicted from contigs of the co-assembly (and therefore MAGs) and (iv) coverage values were calculated for each gene of interest as the number of reads that mapped to each gene divided by the number of bases. Genes and transcripts coverage values were normalized them by the coverage values of the housekeeping gene *gyrB*. On average 26,954,264 DNA sequences and 11.396.571 RNA sequences were mapped onto the assembly (Datasheet 1B). The raw data are accessible at NCBI (SRA accession: PRJNA627699).

### Statistics

Bivariate correlations were calculated between environmental and biological parameters as Spearman rank using a 2-tailed test of significance using the function *rcorr* from the R package Hmisc (v3.6.1) [67] and visualized using the function *corrplot* from the R package corrplot (v0.90) [68].

## Acknowledgements

This work was funded by the Swedish research council Formas (grant 2018-01031), The Carl Trygger Foundation for Scientific Research (grant CTS 18:41), Kempestiftelserna (grants SMK-1753 and SMK-1243), the EMFF-Blue Economy project MER-CLUB (grant 863584), and Marie Curie Individual Fellowship (H2020-MSCA-IF-2016; project-749645). We thank the Swedish Meteorological and Hydrological Institute for letting us participate on their research cruises with the R/V Aranda and we thank the ship crew. We thank the SNP&SEQ Technology Platform in Uppsala. that is a part of the National Genomics Infrastructure (NGI) Sweden and Science for Life Laboratory for the sequencing. The SNP&SEQ Platform is also supported by the Swedish Research Council and the Knut and Alice Wallenberg Foundation. The computations were performed on resources provided by SNIC through Uppsala Multidisciplinary Center for Advanced Computational Science (UPPMAX) using the compute project SNIC 2021/5-53.

## Supporting Information

This file includes descriptions about the Hg speciation and PLS models used, description of molecular analysis and bioinformatics methods, the composition and metabolism of the prokaryotic community, the correlation plots, the phylogenetic trees visualization, and the DOM composition of the studied water samples.

## Datasheet 1

This file includes information related to different parameters collected or measured in this work from samples collected at stations BY32 and BY15 (A) Data related to water depths profiles of ancillary parameters (temperatures, salinity, oxygen, hydrogen sulfide), HgT, MeHg concentrations, MeHg/HgT molar ratio (%) and Hg^II^ methylation rate constant (*k*_meth_) and *hgc* gene and transcript normalized abundances (B) DNA and RNA concentrations measured by Qubit and Nanodrop, number of raw and trimmed reads. (C) Data related to each detect hgc genes, their amino acid sequences, abundance metrics, taxonomic identification and bins id. (D) Abundance data of the 39 functional genes extracted from the metagenomes and metatranscriptomes (E) Data related to the reconstruction of good-quality MAGs from metagenomes (F) Coverage values of the 39 functional genes retrieved in the 10 hgc-carrying MAGs normalized by *gyrB* coverage values (G) Coverage values of *gyrB* and all functional genes retrieved in the 10 *hgc*-carrying MAGs.

## Supplementary information

### Speciation model

The chemical speciation of Hg^II^ and MeHg was determined by thermodynamic modeling using the Solgaswater (WinSGW) software (Karlsson & Lindgren, 2013). We adapted a previously developed model (Soerensen *et al*. 2018, Liem-Nguyen *et al*. 2017) and the model matrix is given in Table S1 and the input concentrations and modeling output for Hg^II^ for each sample is given in Table S2. The pH and Cl^-^ concentration were fixed at 7.3 and 90 mM, respectively for all the samples. The concentration of thiols associated with DOM (DOM-RSH) were calculated as 0.15% mass fraction DOM-RSH of the DOC concentration (Skyllberg *et al*. 2006). The DOC concentration was measured at all 6 depths at BY32 and at 6 of the 12 depths at BY15. A second order polynomial function was fitted to calculate a DOC concentration at the remaining depths. MeHg species were included in the model for completion but are not further discussed.

**Table S1.** Reactions and logarithmic thermodynamic formation constants (log K_f_) for Hg^II^ and MeHg species included in the speciation models.

| Species | Log $K_f$ | Components | | | | | |
| --- | --- | --- | --- | --- | --- | --- | --- |
| | | $\text{H}^+$ | $\text{Cl}^-$ | DOM-RS$^-$ | $\text{H}_2\text{S}$ | $\text{Hg}^{2+}$ | MeHg$^+$ |
| $\text{OH}^-$ | -13.7 | -1 | | | | | |
| $\text{HS}^-$ | -7 | -1 | | | 1 | | |
| DOM-RSH | 9 | 1 |  | 1 |  |  |  |
| $\text{HgOH}^+$ | -3.4 | -1 | | | | 1 | |
| $\text{Hg}(\text{OH})_2$ | -6.2 | -2 | | | | 1 | |
| $\text{HgCl}^+$ | 7.1 | | 1 | | | 1 | |
| $\text{HgCl}_2$ | 13.8 | | 2 | | | 1 | |
| $\text{HgCl}_3^-$ | 14.7 | | 3 | | | 1 | |
| $\text{HgCl}_4^-$ | 15.4 | | 4 | | | 1 | |
| $\text{HgOHCl}$ | 7.8 | -1 | 1 | | | 1 | |
| $\text{Hg}(\text{DOM-RS})_2$ | 41 | | | 2 | | 1 | |
| $\text{HgSH}^+$ | 13.72 | -1 | | | 1 | 1 | |
| $\text{HgS}_2\text{H}^-$ | 18.1 | -3 | | | 2 | 1 | |
| $\text{HgS}_2^{2-}$ | 9 | -4 | | | 2 | 1 | |
| $\text{Hg}(\text{SH})_2$ | 24.6 | -2 | | | 2 | 1 | |
| $\text{HgClSH}$ | 18.89 | -1 | 1 | | 1 | 1 | |
| $\text{HgOHSH}$ | 9.42 | -2 | | | 1 | 1 | |
| $\text{HgS}(\text{s})$ | 30.3 | -2 | | | 1 | 1 | |
| MeHgOH | -4.5 | -1 |  |  |  |  | 1 |
| MeHgCl | 5.4 |  | 1 |  |  |  | 1 |
| MeHg(DOM-RS) | 16.5 |  |  | 1 |  |  | 1 |
| MeHgSH | 7.62 | -1 |  |  | 1 |  | 1 |
| MeHgS$^-$ | 0.12 | -2 | | | 1 | | 1 |
| $\text{S}(\text{MeHg})_2$ | 16.42 | -2 | | | 1 | | 2 |

### Molecular analysis and bioinformatics

Trimmomatic v0.36 was used to quality trim the data with the following parameters: LEADING:5 TRAILING:5 MINLEN:36 ILLUMINACLIP:TruSeq3-PE-2.fa:2:30:10:2 (Bolger *et al*. 2014). The final quality was validated using FastQC v0.11.5 (Andrews 2010). The metagenome assembly of the DNA samples, generated using the assembler MEGAHIT v1.1.2 (Li *et al*. 2015) with default settings, yielded 3,859,129 contigs. The DNA and RNA reads were mapped against the contigs with the function bowtie2-build and bowtie2 from the software bowtie2 (Langmead & Salzberg 2012) and the resulting .sam files were converted to .bam files using the function *samtools* in the software samtools v1.9 (Li *et al*. 2009). This was followed by annotation of the contigs with the *prokka* function in Prokka v1.12 (Seemann 2014) that uses Prodigal v2.6.3 (Hyatt *et al*. 2010) for prokaryotic gene prediction and BLAST v2.6.0+ (Altschul *et al*. 1990) for sequence alignment of translated nucleotides. Prokka was run with a metagenome setup against the UniProtKB/Swiss-Prot protein database (Release18Feb2020) with the following settings: --proteins uniprot_sprot.fasta --metagenome. The .bam files and the prodigal output .gff file were used to estimate sequence counts by using the function *featureCounts* in the software Subread v1.5.1 (Liao *et al*. 2014).

Bins were generated using the function *metabat2* of the software Metabat2 v2.12.1 (Kang *et al*. 2019) (Datasheet 1E). The function *checkM* from the software checkM v1.1.2 (Parks *et al*. 2015) was used to estimate the completeness and redundancy of each bin with the following settings: -- taxonomy_wf life Prokaryote. The software GTDB-Tk v0.3.2 (Chaumeil *et al*. 2019) was used with the following settings: -- classify_wf to taxonomically classify each bin. To annotate the contigs and calculate their coverage in each bin, the functions *prokka* (Prokka v1.12 (Seemann 2014) and *jgi_summarize_bam_contig_depths* (Metabat2 2.12.1 Kang *et al*. 2019) were used. Out of the 280 bins generated in this study (Datasheet 1E), 50, 77 and 121 were respectively defined as high-quality draft (>90% complete, <5% contamination), medium-quality draft (>50% complete, <10% contamination) or low-quality draft (<50% complete, <10% contamination) MAGs respectively. Finally, 32 bins had no reported completeness and contamination values and were not even classified. In the present work, we referred to MAGs only for high-quality and medium-quality MAGs, the low-quality MAGs being not considered as true signals. Altogether 127 good-quality MAGs were reconstructed from the 12 metagenomes (Fig. S1B).

The metabolic capacity of overall microbial community and of specific MAGs was evaluated using the hidden Markov models (HMMs) of Pfam (Finn et al. 2010) and TIGRFAM (Selengut et al. 2007) databases for 39 functional genes (Datasheet 1D) and applying the function *hmmsearch* from the software hmmer v3.2.1 (Finn et al. 2011). Verified hits of the 39 functional genes were detected using the trusted cut-off provided in each HMM file. For each sample and each functional gene, coverage values (number of reads per base) (see Datasheet 1D). The housekeeping gene *gyrB* was detected using the HMM profiles TIGR01059.hmm and applying the trusted cut-off provided in HMM files. The overall community composition was evaluated using the *kraken2* function from kraken2 (Wood et al. 2019) with default settings (v2.0.8) for the taxonomic classification of the sequences obtained in the metagenomes and metatranscriptomes.

### Detection, taxonomic identification and normalization of *hgc* genes and transcript

We first looked for *hgc* gene homologs in the 5,433,642 predicted protein coding genes with the function *hmmsearch* from hmmer (Finn et al. 2011) software (v3.2.1) and using the HMM profiles provided by Hg-MATE database (v1.01142021, doi:10.25573/serc.13105370) built from multiple sequences alignments of *hgcA* and *hgcB* concatenated amino acid sequences. We considered genes with E-values < 10^-3^ as significant hits resulting in 5,174 hits. To identify which of the hit genes truly correspond to *hgcA* and *hgcB* genes, we used the knowledge from the seminal paper of Parks et al. [8] that described unique motifs from *hgcA* (NVWCA(A/G/S)GK) and *hgcB* genes (C(M/I)EC(G/S)(A/G)C) and performed a manual check of the presence of *hgcA* and *hgcB* genes in our dataset resulting in 126 confirmed *hgc* genes, 77 *hgcA* and 49 *hgcB* genes being identified. In this work, we called the detected genes *hgc* genes; however, because their role in Hg methylation could not be verified solely based on environmental genomics data, they are actually defined as putative *hgc* genes. In some cases, *hgcA* and *hgcB* genes were found side-by-side on the same contig. Overall, we detected 23 *hgcAB* gene pairs and 54 *hgcA* genes. The *hgc* genes reads that were detected in sampling and extraction control metagenomes were considered as absent from samples in they account for less than 6 reads as it is the maximum number of reads of *hgc* genes found in control metagenomes. Coverage values of each hgc gene and transcripts were calculated as the number of reads mapped to each gene divided by the number of bases (reads/bp). Genes and transcripts coverage values were normalized by dividing them with the coverage values of the housekeeping gene *gyrB*.

To taxonomically identify hgc-carrying microorganisms detected in BY32 and BY15 samples, we coupled information obtained from *hgc* phylogeny and metagenome-assembled genomes (MAG) that carried *hgc* genes. For *hgc* phylogeny, we used the reference package ‘hgcA’ and ‘hgcAB’ from the recent database Hg-MATE v1.01142021 (Gionfriddo et al. 2021) that compiled a total of 1020 *hgc* sequences from publicly available isolate genomes (n =204), single-cell genomes (n=29) and metagenome-assembled genomes (n=787). Briefly, amino acid sequences from gene previously identified as hgcA gene were (i) compiled in a FASTA file, (ii) aligned to Stockholm formatted alignment of HgcA amino acid sequences from the reference package with the function *hmmalign* from hmmer software v3.2.1 (Finn *et al*. 2011), (iii) placed onto the HgcA reference tree with the function *pplacer* and (iv) classified using the functions *rppr* and *guppy_classify* from the program pplacer (Masten *et al*. 2010). For more details, see in the README.txt of Hg-MATE v1.01142021 (Gionfriddo *et al*. 2021). Among the 127 good-quality MAGs obtained in the present study, ten featured *hgc* genes in their genomes hereafter described as *hgc*-carrying MAGs (Fig. S1, Datasheet 1E). Eight of the *hgc*-carrying MAGs carried *hgc* gene pairs while two had only *hgcA* genes in their genomes (*bin036* ad *bin143*). One MAG, *bin225*, included two *hgc* gene pairs and the strain heterogeneity of this MAG was 100 % (Datasheet 1E) indicating that it likely represents multiple genomes. The 10 *hgc*-carrying MAGs were taxonomically identified as members of Desulfobacterota (1 desulfobacteral *Desulfobacula*, 1 desulfobulbal *Desulforhopalus*, 1 desulfatiglandal NaphS2), PVC superphylum (2 Planctomycetes, 1 Kiritimatiellae, 1 Lentisphaerae), and three other microbial phyla (Chloroflexota, Myxococcota and AABM12-125-24) (Fig. S1). The remaining *hgc* genes were affiliated to Desulfobacterota (n=43), PVC superphylum (n=27), Chloroflexota (n=9) and various bacterial and archaeal lineages (3 Bacteroidetes, 1 Spirochaetes, 1 Firmicutes, 1 Elusimicrobia, 1 Thermoplasmatota) and unidentified microorganisms (n=20) (Datasheet 1C).

### The composition and metabolism of the prokaryotic community from the Central Baltic Sea is linked to redox conditions of its oxygen-deficient water column

The exploration of the metagenomes and metatranscriptomes obtained from the Landsort Deep and the Gotland Deep water columns allowed us to explore the composition and activity of their resident prokaryotic communities. Overall, a total of 127 good-quality metagenome-assembled genomes (MAGs) were recovered from the water column of the two studied stations (BY32 and BY15) (Fig. S1C, Datasheet 1E). High numbers of MAGs were identified as Actinobacteria (n=21), Gammaproteobacteria (n=12), Verrucomicrobia (n=12) and Planctomycetes (n=11). Additionally, the (normalized) gene abundance of these 56 MAGs accounted for 60 % of the overall prokaryotic community although they accounted for lower transcript abundance (27 %). At both stations, transcripts from Synechococcales (Cyanobacteria) and photosynthesis genes were predominant in the normoxic zone (25 m) (Fig. S1A & S1B) in agreement with previous knowledge about the repeated occurrences of *Synechococcus* in the water from the Central Baltic Sea (Stal *et al*. 2003). The underlying redox transition zone exhibited increased amounts of transcripts from Thermoproteota (Fig. S1A) and more specifically the MAG *bin123 (Nitrosopumilus* sp., Nitrososphaerales) (Datasheet 1E). Consistently, members of this obligately aerobic nitrifying archaeal genus have been previously reported in oxygen-deficient waters in the central Baltic Sea (Labrenz *et al*. 2010, Feike *et al*. 2011). Additionally, transcripts from Campylobacterota were detected as highly abundant in the deepest euxinic zones (Fig. S1A). These transcripts were expressed by *Sulfurimonas* sp., (3 dominant MAGs, Datasheet 1E), which are bacteria known to perform denitrification and sulfide oxidation in oxygen-deficient waters including in the Baltic Sea, e.g., (Grote *et al*. 2008, Bergen *et al*. 2018, Beier *et al*. 2019). Their occurrence was coupled with increased transcripts of denitrification and sulfide oxidation genes peaking in both redox transition zones (75 and 80 m at BY32 and BY15 stations, respectively). Although these metabolic functional genes were less abundant in deeper samples, Campylobacterales transcripts remain abundant in underlying water layers supporting knowledge that these bacteria can potentially exhibit metabolic versatility (Grote *et al*. 2008, Beier *et al*. 2019). Finally, Desulfobacterota, known to include numerous sulfate-reducing bacteria (Muyzer & Stams 2008), were more abundant in the euxinic zone (Fig. S1A) in association with increased abundance of transcripts from sulfate reduction genes (Fig. S1B) and hydrogen sulfide concentrations (Fig. 2) in line with previous knowledge about the role of sulfate-reducing Desulfobacterota in the metabolism of oxygen-deficient water columns Canfield *et al*. 2010, Van Vliet *et al*. 2020).

**Figure S1.**
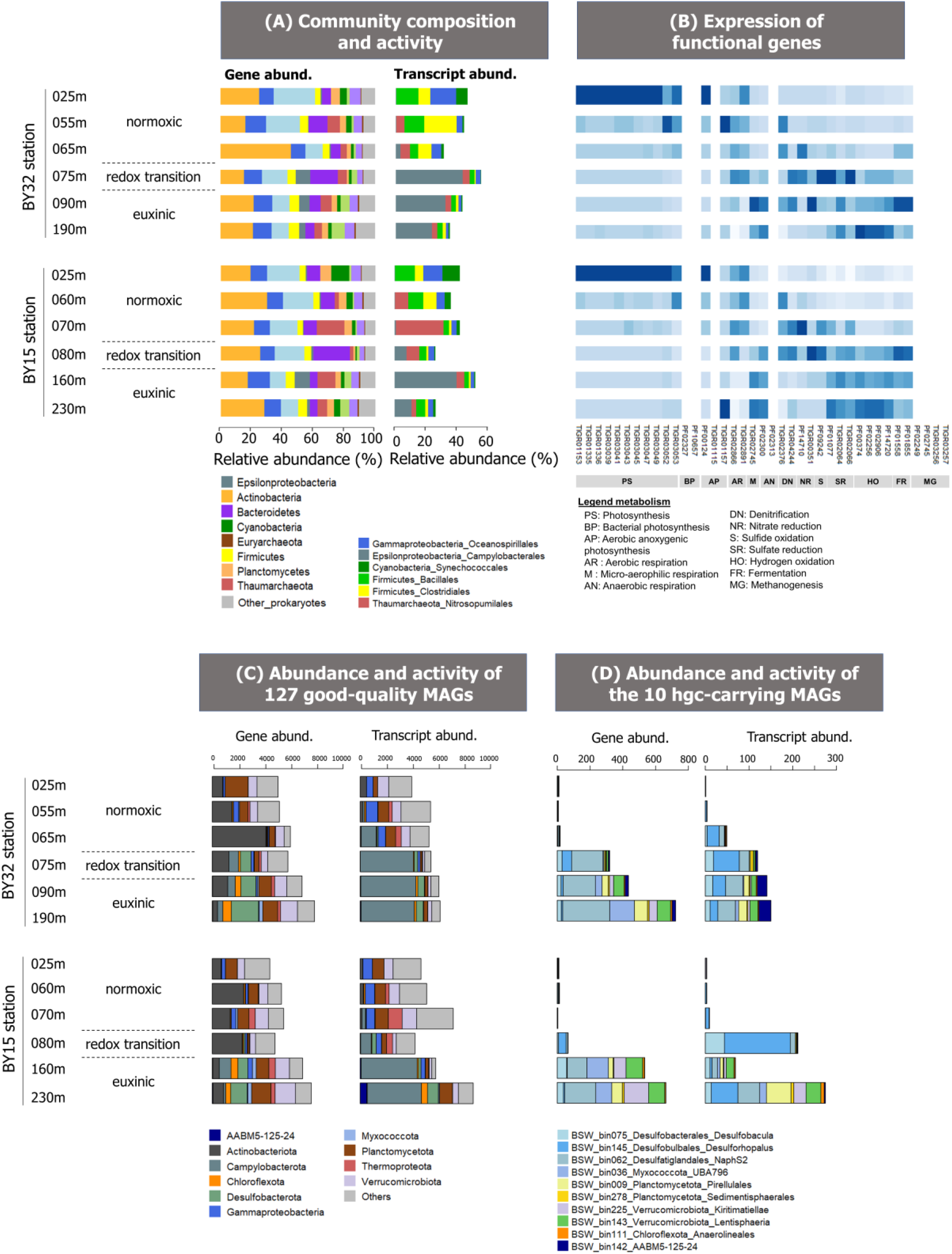
(A) Community composition and activity (B) Heatmap showing the z-scored expression level of each of the 39 functional genes analyzed in the present study covering a broad spectrum of microbial metabolism (e.g., photosynthesis, sulfur cycle and fermentation) (C) Representation of MAGs in terms of normalized total gene and transcript coverage values (D) Representation of the 10 *hgc*-carrying MAGs in terms of normalized total gene and transcript coverage values. Color codes correspond to the taxonomic identification of each microbial group.

**Figure S2:**
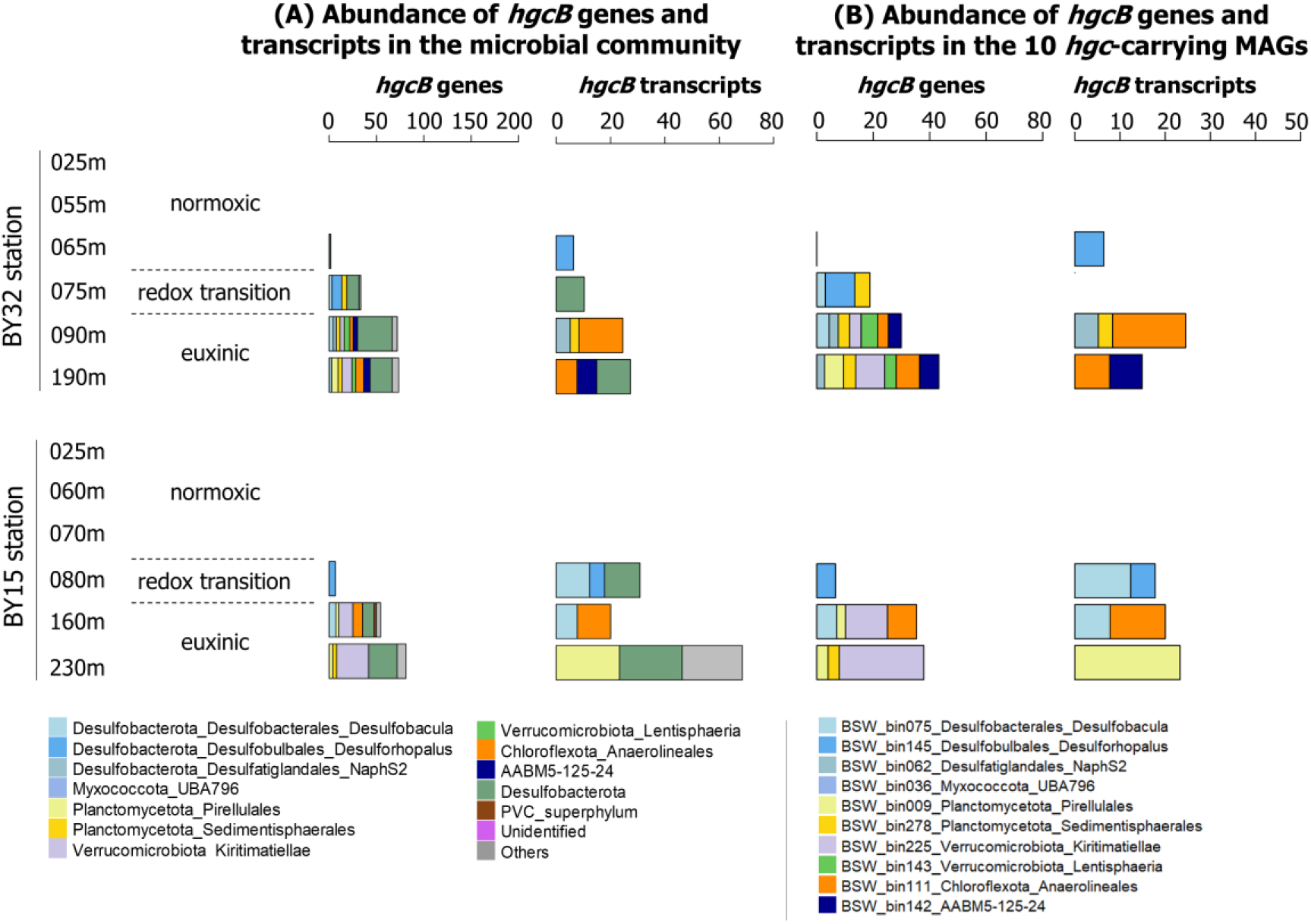
Distribution of *hgcB* genes in terms of normalized abundance (counts) of both total genes and transcripts for all *hgcB* genes detected in the water metagenomes (A) and the 10 *hgc*-carrying MAGs (B). Color codes correspond to the taxonomic identification of each microbial group, see bottom panels for correspondence.

**Table S2.**
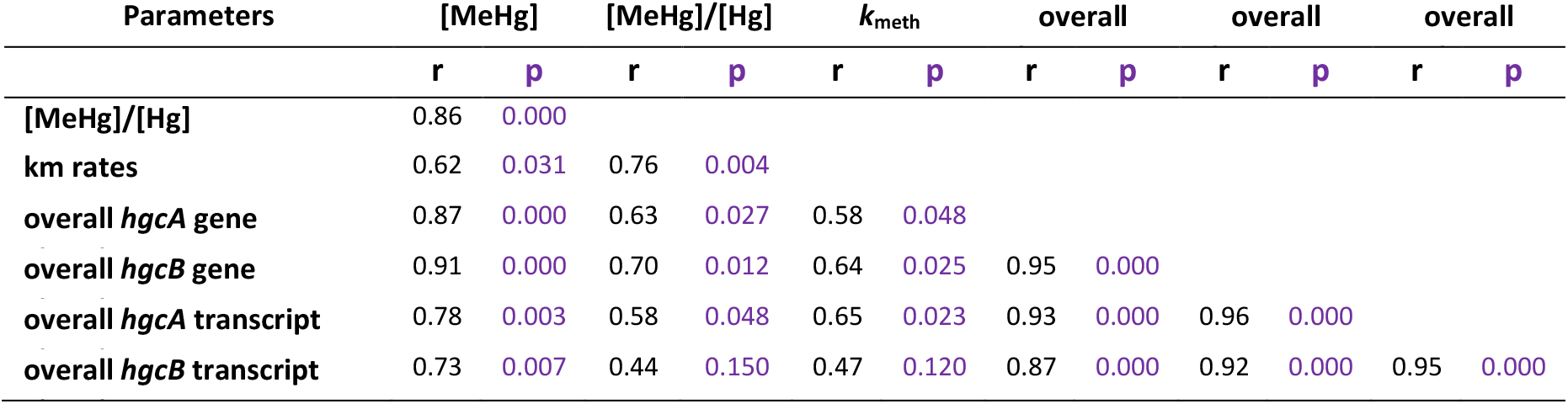
Spearman rank correlations calculated between selected environmental and biological parameters from the overall prokaryotic community. Abundance was calculated as coverage values (reads/bp) normalized by the coverage values of the housekeeping *gyrB*. The r values denote Spearman’s rho rank correlation coefficients, p denotes p-values.

### PLS model

Partial least squares (PLS) projection models were generated in the Simca 15 (Sartorius Stedim Data Analytics AB) software. Separate models (N=12 for each model) were generated with either the *hgcA* gene or transcript abundances as quantitative x-variables. In all models, the concentration of dissolved Hg^II^-sulfide complexes (sum of Hg(SH)2, HgS2H^-^ and HgS2^2-^) were used as a qualitative x-variable (0 or 100%) based on the result of the speciation modeling (Table S2). X-variables were centered and scaled to unit variance. Separate models were generated with *k*_meth_, MeHg concentration or MeHg/HgT molar ratio as y-variable. For each model, two PLS components were fitted. The generated models were thus equivalent to multiple linear regression models with each model consisting of two x-variables and one y-variable. The Simca software and PLS models were used for technical reasons.

**Table S3.** Input concentrations and speciation modeling output (% distribution of major species) for all water samples.

|  |  | Input parameters |  |  |  | Model output |  |  |  |
| --- | --- | --- | --- | --- | --- | --- | --- | --- | --- |
|  | Depth | Hg<sup>II</sup> | MeHg | H<sub>2</sub>S | DOM-RSH | Hg(SH)<sub>2</sub> | HgS<sub>2</sub>H<sup>-</sup> | HgS<sub>2</sub><sup>2-</sup> | Hg(DOM-RS)<sub>2</sub> |
| BY15 | (m) | (fM) | (fM) | (μM) | (μM) | (%) | (%) | (%) | (%) |
| Normoxic | 5 | 851 | 49 | 0 | 0.174 | 0 | 0 | 0 | 100 |
|  | 25 | 401 | 49 | 0 | 0.164 | 0 | 0 | 0 | 100 |
|  | 40 | 701 | 49 | 0 | 0.158 | 0 | 0 | 0 | 100 |
|  | 60 | 651 | 49 | 0 | 0.15 | 0 | 0 | 0 | 100 |
|  | 65 | 419 | 81 | 0 | 0.148 | 0 | 0 | 0 | 100 |
|  | 70 | 801 | 49 | 0 | 0.147 | 0 | 0 | 0 | 100 |
|  | 75 | 501 | 49 | 0 | 0.145 | 0 | 0 | 0 | 100 |
| Transition | 80 | 951 | 49 | 0 | 0.143 | 0 | 0 | 0 | 100 |
|  | 130 | 973 | 227 | 0 | 0.133 | 0 | 0 | 0 | 100 |
| Euxinic | 160 | 2250 | 950 | 22 | 0.127 | 10.1 | 86.5 | 3.4 | <0.001 |
|  | 195 | 446 | 1454 | 44 | 0.125 | 10.1 | 86.5 | 3.4 | <0.001 |
|  | 230 | 790 | 1510 | 100 | 0.126 | 10.1 | 86.5 | 3.4 | <0.001 |
| BY32 |  |  |  |  |  |  |  |  |  |
| Normoxic | 25 | 351 | 49 | 0 | 0.173 | 0 | 0 | 0 | 100 |
|  | 55 | 451 | 49 | 0 | 0.159 | 0 | 0 | 0 | 100 |
|  | 65 | 501 | 49 | 0 | 0.155 | 0 | 0 | 0 | 100 |
| Transition | 75 | 691 | 859 | 0 | 0.152 | 0 | 0 | 0 | 100 |
| Euxinic | 90 | 275 | 1275 | 30 | 0.148 | 10.1 | 86.5 | 3.4 | <0.001 |
|  | 190 | 1033 | 867 | 41 | 0.141 | 10.1 | 86.5 | 3.4 | <0.001 |

**Figure S3.**
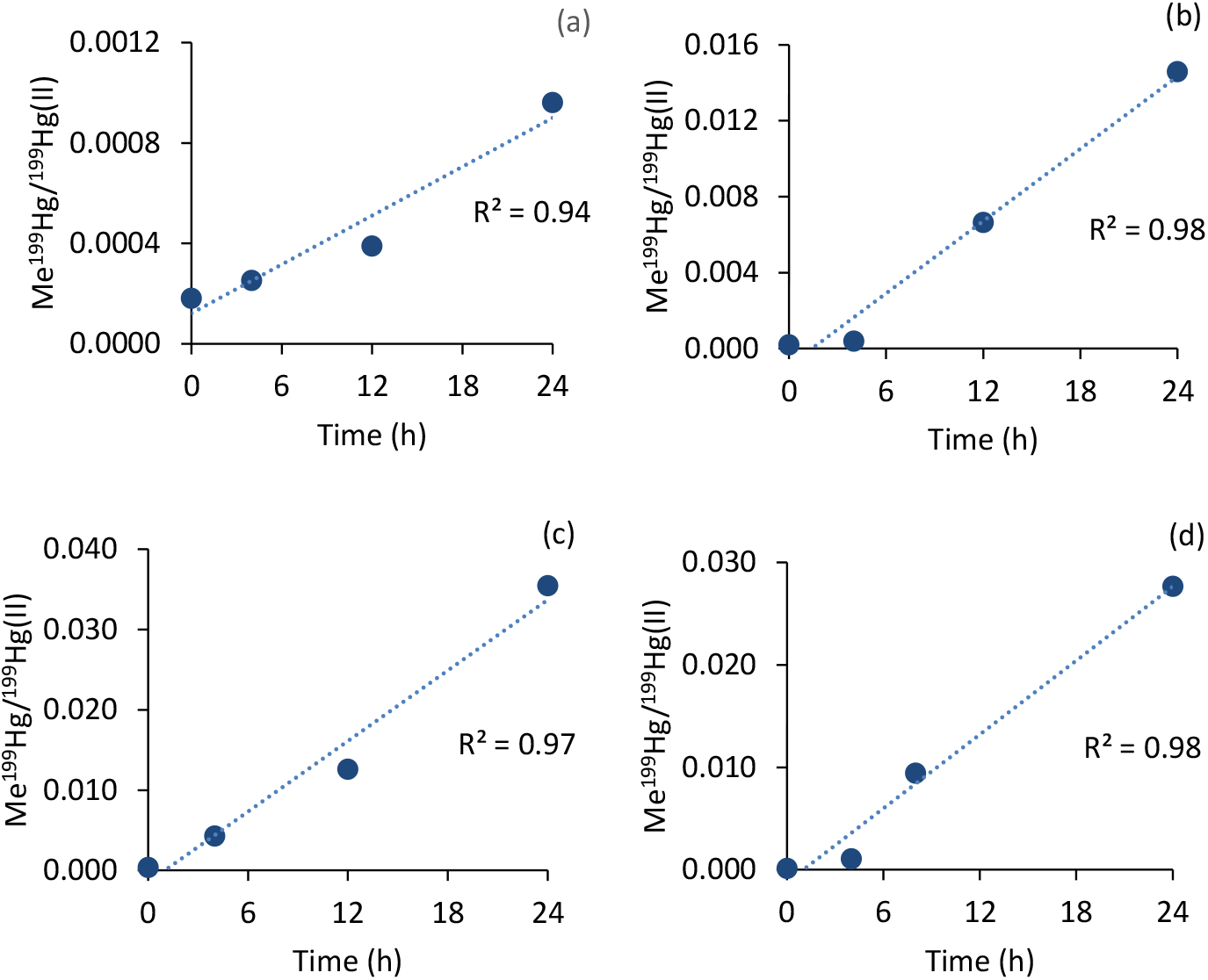
Formation of Me^199^Hg from added ^199^Hg(II) (expressed as Me^199^Hg/^199^Hg(II) molar ratio) over time in water sample incubation experiments for sample with a significant Hg(II) methylation rate constant (*k*_meth_), i.e. samples (a) BY32 75m, (b) BY32 90m, (c) BY32 190m and (d) BY15 230 m.

### The DOM composition of the redox gradient from the Central Baltic Sea is stable

In this first DOM composition data interpretation, there was one clear outlier (Bal-BY32-25m). The pair of Suwannee River Fulvic Acid reference material analyses gave a dissimilarity of 2.9%, indicating high method precision. The rest of the Baltic samples were highly similar to each other, with dissimilarity values generally <10%. Fig. S7 shows a heatmap of the Baltic samples alone, without the outlier. Overall, the molecular composition of DOM was found to be very similar at both stations and at all water depths. In particular, no pattern was found with depth (e.g., above and below the redox boundaries), indicating that compositional differences in ionizable DOM molecules do not explain differences in Hg species abundances or formation potential. Rather, our data support that these parameters are controlled by the presence of inorganic sulfide species and *hgc*-carrying microorganisms.

**Figure S4.**
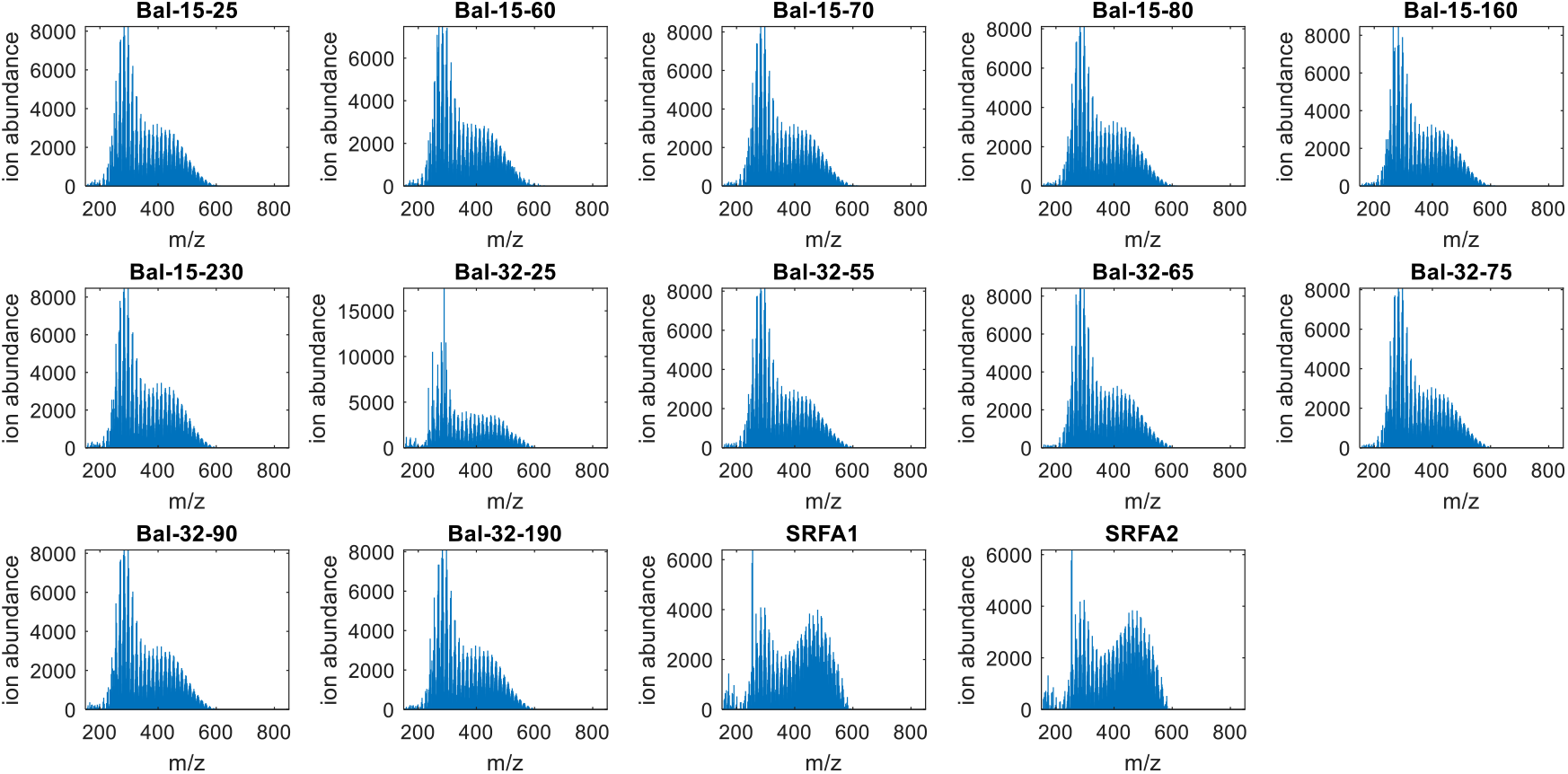
Mass spectra of the extracted DOM samples.

**Figure S5.**
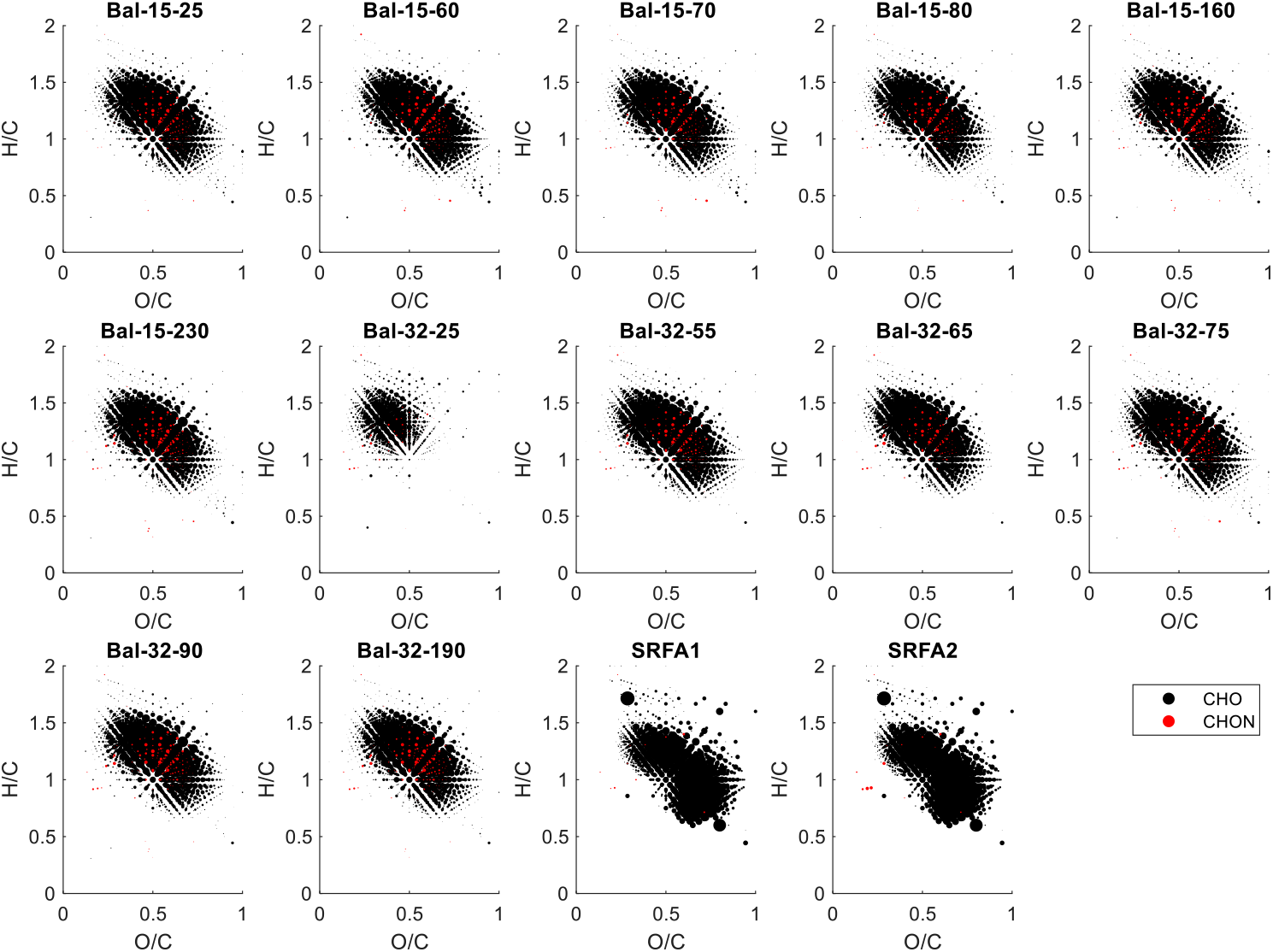
van Krevelen diagrams (H/C vs O/C ratio) of assigned formulas for the extracted DOM samples. Point size indicates summed peak intensity.

**Figure S6.**
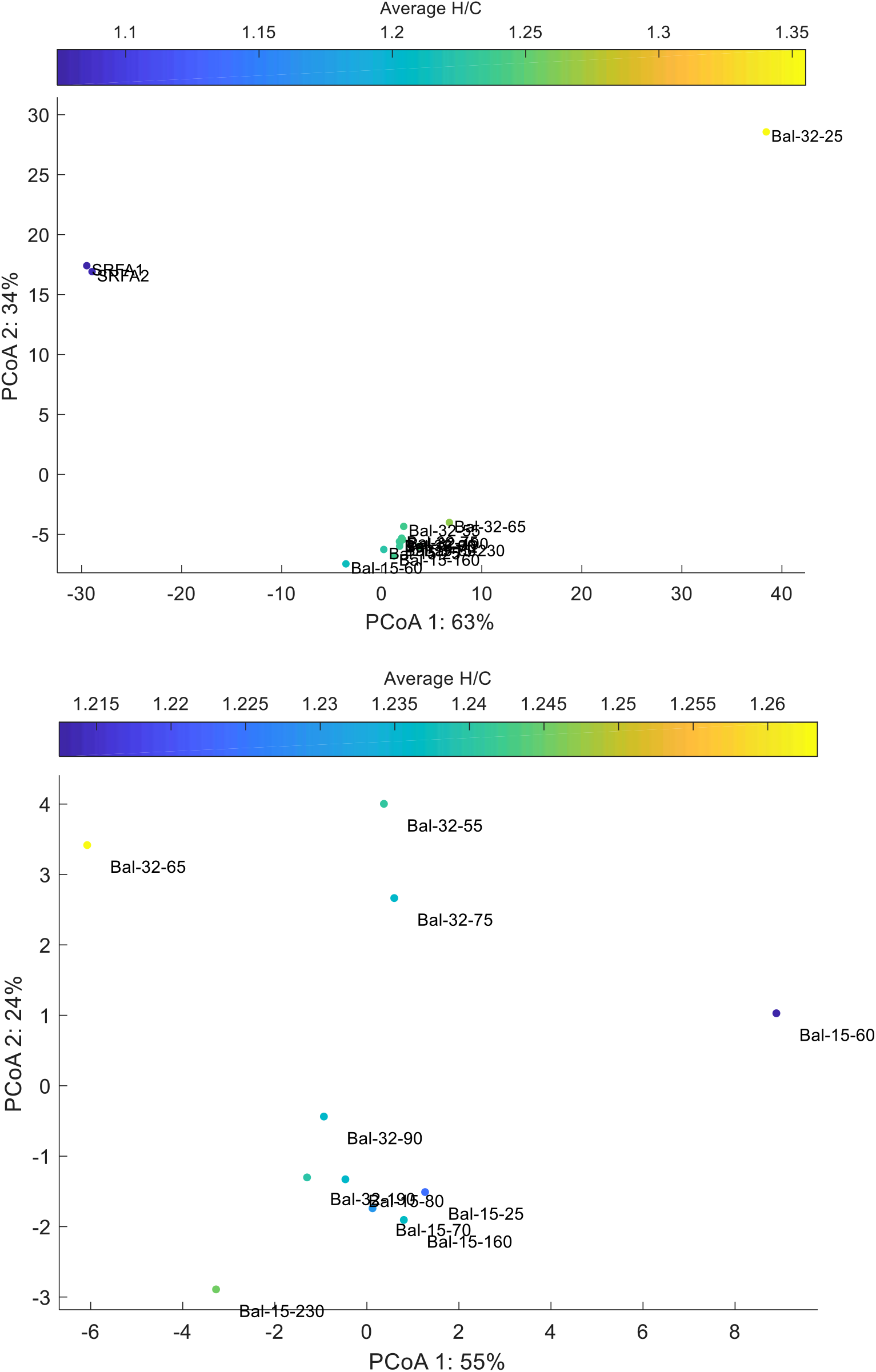
The top figure shows the principal coordinate analysis of Bray Curtis Dissimilarities for all sample, indicating high dissimilarity between the majority of samples and site 32 at 25m, and also the reference mixture of SRFA. The bottom figure shows the principal coordinate diagram of non-outlying samples.

**Figure S7.**
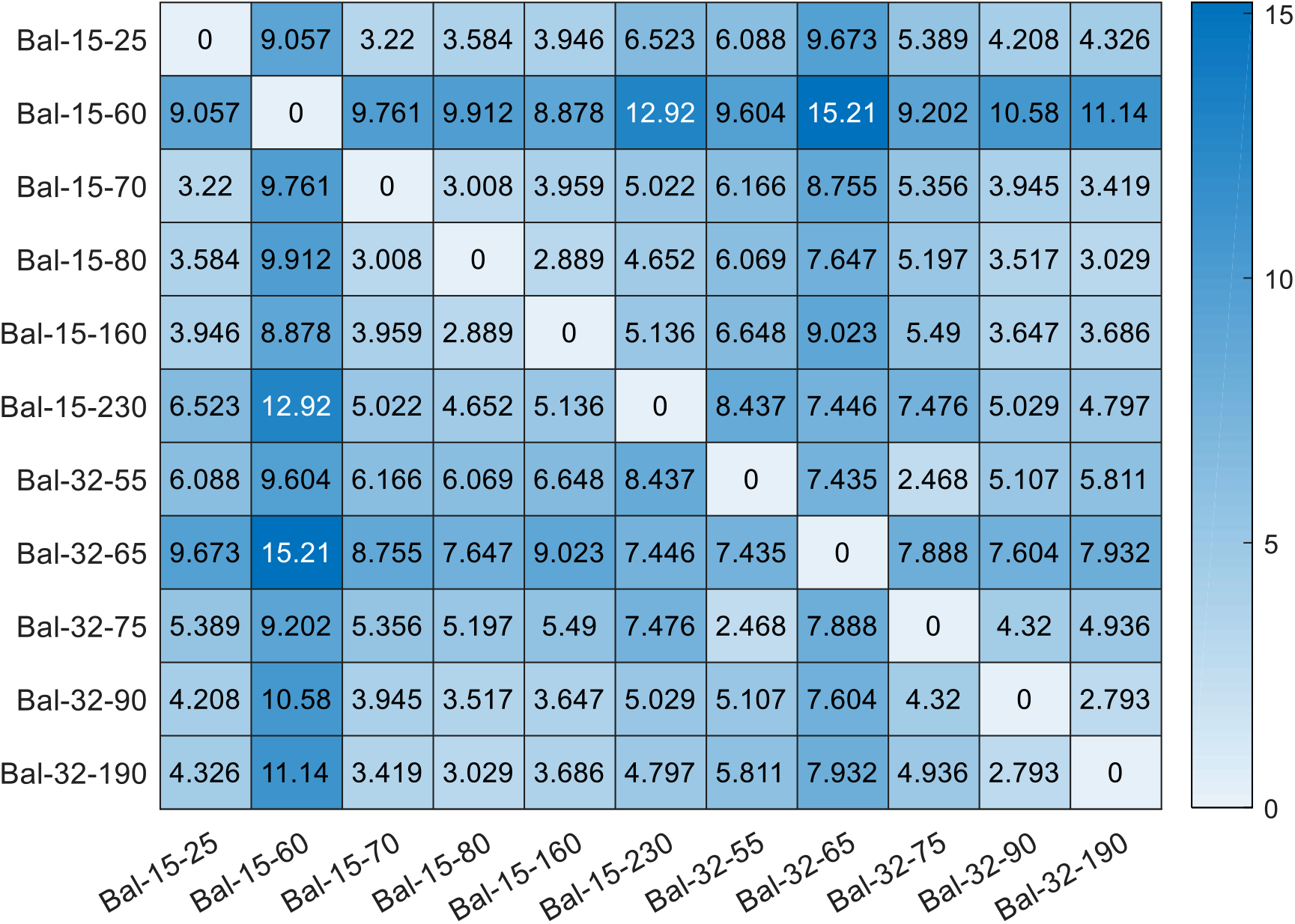
Heatmap of Bray Curtis Dissimilarities for the molecular composition of the extracted DOM samples.

## References

1. Driscoll CT, Mason RP, Chan HM, Jacob DJ, Pirrone N. Mercury as a Global Pollutant: Sources, Pathways, and Effects. Environ Sci Technol 2013; 47: 4967–4983.

2. Selin NE. A proposed global metric to aid mercury pollution policy. Science 2018; 360: 607–609.

3. Munthe J, Bodaly RA, Branfireun BA, Driscoll CT, Gilmour CC, Harris R, et al. Recovery of Mercury-Contaminated Fisheries. Ambio 2007; 36: 33–44.

4. Krabbenhoft DP, Sunderland EM. Global Change and Mercury. Science 2013; 341: 1457–1458.

5. Hsu-Kim H, Eckley CS, Achá D, Feng X, Gilmour CC, Jonsson S, et al. Challenges and opportunities for managing aquatic mercury pollution in altered landscapes. Ambio 2018; 47: 141–169.

6. Kidd K, Clayden M, Jardine T. Bioaccumulation and Biomagnification of Mercury through Food Webs. Environmental Chemistry and Toxicology of Mercury. 2011. John Wiley & Sons, Inc., Hoboken, NJ, USA, pp 453–499.

7. Doney SC. The growing human footprint on coastal and open-ocean biogeochemistry. Science 2010; 328: 1512–6.

8. Parks JM, Johs A, Podar M, Bridou R, Hurt RA, Smith SD, et al. The genetic basis for bacterial mercury methylation. Science 2013; 339: 1332–1335.

9. Bravo AG, Cosio C. Biotic formation of methylmercury: A bio–physico–chemical conundrum. Limnol Oceanogr 2020; 65: 1010–1027.

10. Compeau GC, Bartha R. Sulfate-reducing bacteria: Principal methylators of Mercury in Anoxic Estuarine Sediment. Appl Environ Microbiol 1985; 50: 498–502.

11. Fleming EJ, Mack EE, Green PG, Nelson DC. Mercury methylation from unexpected sources: molybdate-inhibited freshwater sediments and an iron-reducing bacterium. Appl Environ Microbiol 2006; 72: 457–64.

12. Bravo AG, Zopfi J, Buck M, Xu J, Bertilsson S, Schaefer JK, et al. Geobacteraceae are important members of mercury-methylating microbial communities of sediments impacted by waste water releases. ISME J 2018; 12: 802–812.

13. Peterson BD, McDaniel EA, Schmidt AG, Lepak RF, Janssen SE, Tran PQ, et al. Mercury Methylation Genes Identified across Diverse Anaerobic Microbial Guilds in a Eutrophic Sulfate-Enriched Lake. Environ Sci Technol 2020; 54: 15840–15851.

14. Yu RQ, Reinfelder JR, Hines ME, Barkay T. Syntrophic pathways for microbial mercury methylation. ISME J 2018; 12: 1826–1835.

15. Goñi-Urriza M, Corsellis Y, Lanceleur L, Tessier E, Gury J, Monperrus M, et al. Relationships between bacterial energetic metabolism, mercury methylation potential, and hgcA/hgcB gene expression in Desulfovibrio dechloroacetivorans BerOc1. Environ Sci Pollut Res 2015; 22: 13764–13771.

16. Bravo AG, Kothawala DN, Attermeyer K, Tessier E, Bodmer P, Ledesma JLJ, et al. The interplay between total mercury, methylmercury and dissolved organic matter in fluvial systems: A latitudinal study across Europe. Water Res 2018; 144: 172–182.

17. Christensen GA, Gionfriddo CM, King AJ, Moberly JG, Miller CL, Somenahally AC, et al. Determining the Reliability of Measuring Mercury Cycling Gene Abundance with Correlations with Mercury and Methylmercury Concentrations. Environ Sci Technol 2019; 53: 8649–8663.

18. Conley DJ, Carstensen J, Aigars J, Axe P, Bonsdorff E, Eremina T, et al. Hypoxia Is Increasing in the Coastal Zone of the Baltic Sea. Environ Sci Technol 2011; 45: 6777–6783.

19. Breitburg D, Levin LA, Oschlies A, Grégoire M, Chavez FP, Conley DJ, et al. Declining oxygen in the global ocean and coastal waters. Science 2018; 359: 1–11.

20. Carstensen J, Andersen JH, Gustafsson BG, Conley DJ. Deoxygenation of the Baltic Sea during the last century. Proc Natl Acad Sci U S A 2014; 111: 5628–33.

21. Hansson M, Viktorsson L, Andersson L. Oxygen Survey in the Baltic Sea 2019 - Extent of Anoxia and Hypoxia, 1960-2019. 2020. [Swedish Meteorological and Hydrological Institute].

22. Lamborg CH, Buesseler KO, Valdes J, Bertrand CH, Bidigare R, Manganini S, et al. The flux of bio- and lithogenic material associated with sinking particles in the mesopelagic “twilight zone” of the northwest and North Central Pacific Ocean. Deep Sea Res Part II Top Stud Oceanogr 2008; 55: 1540–1563.

23. Kuss J, Krüger S, Ruickoldt J, Wlost K-P. High-resolution measurements of elemental mercury in surface water for an improved quantitative understanding of the Baltic Sea as a source of atmospheric mercury. Atmos Chem Phys 2018; 18: 4361–4376.

24. Pakhomova S, Braaten H, Yakushev E, Protsenko E. Water Column Distribution of Mercury Species in Permanently Stratified Aqueous Environments. Oceanology 2018; 58: 28–37.

25. Rosati G, Heimbürger LE, Melaku Canu D, Lagane C, Laffont L, Rijkenberg MJA, et al. Mercury in the Black Sea: New Insights From Measurements and Numerical Modeling. Global Biogeochem Cycles 2018; 32: 529–550.

26. Soerensen AL, Schartup AT, Skrobonja A, Bouchet S, Amouroux D, Liem-Nguyen V, et al. Deciphering the Role of Water Column Redoxclines on Methylmercury Cycling Using Speciation Modeling and Observations From the Baltic Sea. Global Biogeochem Cycles 2018; 32: 1498–1513.

27. Feistel R, Nausch G, Wasmund N. State and evolution of the Baltic Sea, 1952-2005: a detailed 50-year survey of meteorology and climate, physics, chemistry, biology, and marine environment. 2008.

28. Elken J, Matthäus W. Physical system description. Assessment of climate change for the Baltic Sea basin, Springer S. 2008.

29. Mason RP, Fitzgerald WF, Hurley J, Hanson AK, Donaghay PL, Sieburth JM. Mercury biogeochemical cycling in a stratified estuary. Limnol Oceanogr 1993; 38: 1227–1241.

30. Drott A, Lambertsson L, Björn E, Skyllberg U. Do potential methylation rates reflect accumulated methyl mercury in contaminated sediments? Environ Sci Technol 2008; 42: 153–8.

31. Chadwick SP, Babiarz CL, Hurley JP, Armstrong DE. Influences of iron, manganese, and dissolved organic carbon on the hypolimnetic cycling of amended mercury. Sci Total Environ 2006; 368: 177–188.

32. Jonsson S, Skyllberg U, Nilsson MB, Westlund P-O, Shchukarev A, Lundberg E, et al. Mercury Methylation Rates for Geochemically Relevant Hg II Species in Sediments. Environ Sci Technol 2012; 46: 11653–11659.

33. Zhang T, Kim B, Levard C, Reinsch BC, Lowry G V., Deshusses MA, et al. Methylation of Mercury by Bacteria Exposed to Dissolved, Nanoparticulate, and Microparticulate Mercuric Sulfides. Environ Sci Technol 2012; 46: 6950–6958.

34. Benoit JM, Gilmour CC, Mason RP, Heyes A. Sulfide Controls on Mercury Speciation and Bioavailability to Methylating Bacteria in Sediment Pore Waters. Environ Sci Technol 1999; 33: 951–957.

35. Drott A, Björn E, Bouchet S, Skyllberg U. Refining Thermodynamic Constants for Mercury(II)-Sulfides in Equilibrium with Metacinnabar at Sub-Micromolar Aqueous Sulfide Concentrations. Environ Sci Technol 2013; 47: 4197–4203.

36. Skyllberg U. Competition among thiols and inorganic sulfides and polysulfides for Hg and MeHg in wetland soils and sediments under suboxic conditions: Illumination of controversies and implications for MeHg net production. J Geophys Res Biogeosciences 2008; 113.

37. Gionfriddo CM, Tate MT, Wick RR, Schultz MB, Zemla A, Thelen MP, et al. Microbial mercury methylation in Antarctic sea ice. Nat Microbiol 2016; 1: 1–12.

38. Villar E, Cabrol L, Heimbürger-Boavida LE. Widespread microbial mercury methylation genes in the global ocean. Environ Microbiol Rep 2020; 12: 277–287.

39. Tada Y, Marumoto K, Takeuchi A. Nitrospina-Like Bacteria Are Potential Mercury Methylators in the Mesopelagic Zone in the East China Sea. Front Microbiol 2020; 11: 1–16.

40. Capo E, Bravo AG, Soerensen AL, Bertilsson S, Pinhassi J, Feng C, et al. Deltaproteobacteria and Spirochaetes-Like Bacteria Are Abundant Putative Mercury Methylators in Oxygen-Deficient Water and Marine Particles in the Baltic Sea. Front Microbiol 2020; 11: 1–11.

41. Capo E, Broman E, Bonaglia S, Bravo AG, Bertilsson S, Soerensen AL, et al. Oxygen-deficient water zones in the Baltic Sea promote uncharacterized Hg methylating microorganisms in underlying sediments. Limnol Oceanogr 2022; 67: 135–146.

42. Lin H, Ascher DB, Myung Y, Lamborg CH, Hallam SJ, Gionfriddo CM, et al. Mercury methylation by metabolically versatile and cosmopolitan marine bacteria. ISME J 2021; 15: 1810–1825.

43. Isaksen MF, Teske A. Desulforhopalus vacuolatus gen. nov., sp. nov., a new moderately psychrophilic sulfate-reducing bacterium with gas vacuoles isolated from a temperate estuary. Arch Microbiol 1996; 166: 160–168.

44. Galushko A, Minz D, Schink B, Widdel F. Anaerobic degradation of naphthalene by a pure culture of a novel type of marine sulphate-reducing bacterium. Environ Microbiol 1999; 1: 415–420.

45. Bolliger C, Schroth MH, Bernasconi SM, Kleikemper J, Zeyer J. Sulfur isotope fractionation during microbial sulfate reduction by toluene-degrading bacteria. Geochim Cosmochim Acta 2001; 65: 3289–3298.

46. Anantharaman K, Hausmann B, Jungbluth SP, Kantor RS, Lavy A, Warren LA, et al. Expanded diversity of microbial groups that shape the dissimilatory sulfur cycle. ISME J 2018; 12: 1715–1728.

47. Yu RQ, Flanders JR, MacK EE, Turner R, Mirza MB, Barkay T. Contribution of coexisting sulfate and iron reducing bacteria to methylmercury production in freshwater river sediments. Environ Sci Technol 2012; 46: 2684–2691.

48. McDaniel E, Peterson B, Stevens S, Tran P, Anantharaman K, McMahon K. Expanded Phylogenetic Diversity and Metabolic Flexibility of. mSystems 2020; 5: 1–21.

49. Vigneron A, Cruaud P, Aubé J, Guyoneaud R, Goñi-Urriza M. Transcriptomic evidence for versatile metabolic activities of mercury cycling microorganisms in brackish microbial mats. npj Biofilms Microbiomes 2021; 7: 1–11.

50. Gilmour CC, Podar M, Bullock AL, Graham AM, Brown SD, Somenahally AC, et al. Mercury methylation by novel microorganisms from new environments. Environ Sci Technol 2013; 47: 11810–11820.

51. Bravo AG, Bouchet S, Tolu J, Björn E, Mateos-Rivera A, Bertilsson S. Molecular composition of organic matter controls methylmercury formation in boreal lakes. Nat Commun 2017; 8: 1–9.

52. Herrero Ortega S, Catalán N, Björn E, Gröntoft H, Hilmarsson TG, Bertilsson S, et al. High methylmercury formation in ponds fueled by fresh humic and algal derived organic matter. Limnol Oceanogr 2018; 63: S44–S53.

53. Hawkes JA, Radoman N, Bergquist J, Wallin MB, Tranvik LJ, Löfgren S. Regional diversity of complex dissolved organic matter across forested hemiboreal headwater streams. Sci Rep 2018; 8: 16060.

54. Podar M, Gilmour CC, Brandt CC, Soren A, Brown SD, Crable BR, et al. Global prevalence and distribution of genes and microorganisms involved in mercury methylation. Sci Adv 2015; 1: 1–13.

55. Qian C, Chen H, Johs A, Lu X, An J, Pierce EM, et al. Quantitative Proteomic Analysis of Biological Processes and Responses of the Bacterium Desulfovibrio desulfuricans ND132 upon Deletion of Its Mercury Methylation Genes. Proteomics 2018; 18: 1700479.

56. Schaefer JK, Kronberg RM, Björn E, Skyllberg U. Anaerobic guilds responsible for mercury methylation in boreal wetlands of varied trophic status serving as either a methylmercury source or sink. Environ Microbiol 2020; 22: 3685–3699.

57. Rocca JD, Hall EK, Lennon JT, Evans SE, Waldrop MP, Cotner JB, et al. Relationships between protein-encoding gene abundance and corresponding process are commonly assumed yet rarely observed. ISME J 2015; 9: 1693–1699.

58. Cooper CJ, Zheng K, Rush KW, Johs A, Sanders BC, Pavlopoulos GA, et al. Structure determination of the HgcAB complex using metagenome sequence data: insights into microbial mercury methylation. Commun Biol 2020; 3: 1–9.

59. Gilmour CC, Elias DA, Kucken AM, Brown SD, Palumbo A V., Schadt CW, et al. Sulfate-reducing bacterium Desulfovibrio desulfuricans ND132 as a model for understanding bacterial mercury methylation. Appl Environ Microbiol 2011; 77: 3938–3951.

60. Hammerschmidt CR, Bowman KL, Tabatchnick MD, Lamborg CH. Storage bottle material and cleaning for determination of total mercury in seawater. Limnol Oceanogr Methods 2011; 9: 426–431.

61. Lambertsson L, Björn E. Validation of a simplified field-adapted procedure for routine determinations of methyl mercury at trace levels in natural water samples using species-specific isotope dilution mass spectrometry. Anal Bioanal Chem 2004; 380: 871–875.

62. Munson KM, Babi D, Lamborg CH. Determination of monomethylmercury from seawater with ascorbic acid-assisted direct ethylation. Limnol Oceanogr Methods 2014; 12: 1–9.

63. Karlsson M & Lindgren J (2012) www.winsgw.se.

64. Liem-Nguyen V, Skyllberg U, Björn E. Thermodynamic Modeling of the Solubility and Chemical Speciation of Mercury and Methylmercury Driven by Organic Thiols and Micromolar Sulfide Concentrations in Boreal Wetland Soils. Environ Sci Technol 2017; 51: 3678–3686.

65. Dittmar T, Koch B, Hertkorn N, Kattner G. A simple and efficient method for the solid-phase extraction of dissolved organic matter (SPE-DOM) from seawater. Limnol Oceanogr Methods 2008; 6: 230–235.

66. Patriarca C, Balderrama A, Može M, Sjöberg PJR, Bergquist J, Tranvik LJ, et al. Investigating the Ionization of Dissolved Organic Matter by Electrospray. Anal Chem 2020; 92: 14210–14218.

67. Harrell F, Harrell M. Package ‘Hmisc’. CRAN 2013; 235.

68. Wei T, Simko V, Levy M, Xie Y, Jin Y, Zemla J. Package ‘corrplot’. Statistician 2017; 56: e24.

## References for Supporting Information

Altschul SF, Gish W, Miller W, Myers EW, Lipman DJ. Basic local alignment search tool. J Mol Biol 1990; 215: 403–410. Andrews S FastQC: a quality control tool for high throughput sequence data

Beier S, Holtermann PL, Numberger D, Schott T, Umlauf L, Jürgens K. A metatranscriptomics-based assessment of small-scale mixing of sulfidic and oxic waters on redoxcline prokaryotic communities. Environ Microbiol 2019; 21: 584–602.

Bergen B, Naumann M, Herlemann DPR, Gräwe U, Labrenz M, Jürgens K. Impact of a Major inflow event on the composition and distribution of bacterioplankton communities in the Baltic Sea. Front Mar Sci 2018; 5: 1–14.

Bolger A, Lohse M, Usadel B. Trimmomatic: a flexible trimmer for Illumina sequence data. Bioinformatics 2014.

Canfield DE, Stewart FJ, Thamdrup B, De Brabandere L, Dalsgaard T, Delong EF, et al. A cryptic sulfur cycle in oxygen-minimum-zone waters off the Chilean coast. Science (80-) 2010; 330: 1375–8.

Chaumeil P-A, Mussig AJ, Hugenholtz P, Parks DH. GTDB-Tk: a toolkit to classify genomes with the Genome Taxonomy Database. Bioinformatics 2019; 36: 1925–1927.

Feike J, Jürgens K, Hollibaugh JT, Krüger S, Jost G, Labrenz M. Measuring unbiased metatranscriptomics in suboxic waters of the central Baltic Sea using a new in situ fixation system. ISME J 2011; 6: 461–470.

Finn RD, Mistry J, Tate J, Coggill P, Heger A, Pollington JE, et al. The Pfam protein families database. Nucleic Acids Res 2010; 38: D211–D222.

Finn RD, Clements J, Eddy SR. HMMER web server: interactive sequence similarity searching. Nucleic Acids Res 2011; 39: W29–W37.

Gionfriddo C, Capo E, Peterson B, Lin H, Jones D, Bravo A, et al. Hg-cycling Microorganisms in Aquatic and Terrestrial Ecosystems Database v1.01142021. https://doi.org/1025573/serc13105370.v1 2021.

Grote J, Jost G, Labrenz M, Herndl GJ, Jürgens K. Epsilonproteobacteria represent the major portion of chemoautotrophic bacteria in sulfidic waters of pelagic redoxclines of the baltic and black seas. Appl Environ Microbiol 2008; 74: 7546–7551.

Hyatt D, Chen G-L, LoCascio PF, Land ML, Larimer FW, Hauser LJ. Prodigal: prokaryotic gene recognition and translation initiation site identification. BMC Bioinformatics 2010; 11: 119.

Kang DD, Li F, Kirton E, Thomas A, Egan R, An H, et al. MetaBAT 2: an adaptive binning algorithm for robust and efficient genome reconstruction from metagenome assemblies. PeerJ 2019; 7: e7359.

Karlsson M & Lindgren J (2012) www.winsgw.se.

Labrenz M, Sintes E, Toetzke F, Zumsteg A, Herndl GJ, Seidler M, et al. Relevance of a crenarchaeotal subcluster related to Candidatus Nitrosopumilus maritimus to ammonia oxidation in the suboxic zone of the central Baltic Sea. ISME J 2010; 4: 1496–1508.

Langmead B, Salzberg SL. Fast gapped-read alignment with Bowtie 2. Nat Methods 2012; 9: 357–359.

Li H, Handsaker B, Wysoker A, Fennell T, Ruan J, Homer N, et al. The Sequence Alignment/Map format and SAMtools. Bioinformatics 2009; 25: 2078–2079.

Li D, Liu C-M, Luo R, Sadakane K, Lam T-W. MEGAHIT: an ultra-fast single-node solution for large and complex metagenomics assembly via succinct de Bruijn graph. Bioinformatics 2015; 31: 1674–1676.

Liao Y, Smyth GK, Shi W. featureCounts: an efficient general purpose program for assigning sequence reads to genomic features. Bioinformatics 2014; 30: 923–930.

Liem-Nguyen V, Skyllberg U, Björn E. Thermodynamic Modeling of the Solubility and Chemical Speciation of Mercury and Methylmercury Driven by Organic Thiols and Micromolar Sulfide Concentrations in Boreal Wetland Soils. Environ Sci Technol 2017; 51: 3678–3686.

Matsen FA, Kodner RB, Armbrust EV. pplacer: linear time maximum-likelihood and Bayesian phylogenetic placement of sequences onto a fixed reference tree. BMC Bioinformatics 2010; 11: 538.

Muyzer G, Stams AJM. The ecology and biotechnology of sulphate-reducing bacteria. Nat Rev Microbiol 2008; 6: 441–454.

Parks DH, Imelfort M, Skennerton CT, Hugenholtz P, Tyson GW. CheckM: assessing the quality of microbial genomes recovered from isolates, single cells, and metagenomes. Genome Res 2015; 25: 1043–55.

Seemann T. Prokka: rapid prokaryotic genome annotation. Bioinformatics 2014.

Selengut JD, Haft DH, Davidsen T, Ganapathy A, Gwinn-Giglio M, Nelson WC, et al. TIGRFAMs and Genome Properties: tools for the assignment of molecular function and biological process in prokaryotic genomes. Nucleic Acids Res 2007; 35: D260–D264.

Skyllberg U, Bloom PR, Qian J, Chung-Min Lin, Bleam§ WF. Complexation of Mercury(II) in Soil Organic Matter: EXAFS Evidence for Linear Two-Coordination with Reduced Sulfur Groups. Environ Sci Technol 2006; 40: 4174–4180.

Soerensen AL, Schartup AT, Skrobonja A, Bouchet S, Amouroux D, Liem-Nguyen V, et al. Deciphering the Role of Water Column Redoxclines on Methylmercury Cycling Using Speciation Modeling and Observations From the Baltic Sea. Global Biogeochem Cycles 2018; 32: 1498–1513.

Stal L, Albertano P, Bergman B, Bröckel K von, Gallon J, Hayes P, et al. BASIC: Baltic Sea cyanobacteria. An investigation of the structure and dynamics of water blooms of cyanobacteria in the Baltic Sea—responses to a changing. Cont Shelf Res 2003; 23: 1695–1714.

van Vliet DM, von Meijenfeldt FAB, Dutilh BE, Villanueva L, Sinninghe Damsté JS, Stams AJM, et al. The bacterial sulfur cycle in expanding dysoxic and euxinic marine waters. Environ Microbiol 2020.

Wood DE, Lu J, Langmead B. Improved metagenomic analysis with Kraken 2. Genome Biol 2019; 20: 257.

